# Uncovering the effects of Müllerian mimicry on the evolution of conspicuousness in colour patterns

**DOI:** 10.1101/620963

**Authors:** Ombeline Sculfort, Ludovic Maisonneuve, Marianne Elias, Thomas G. Aubier, Violaine Llaurens

## Abstract

Variation in the conspicuousness of colour patterns is observed within and among defended prey species. The evolution of conspicuous colour pattern in defended species can be strongly impaired because of increased detectability by predators. Nevertheless, such evolution of the colour pattern can be favoured if changes in conspicuousness result in Müllerian mimicry with other defended prey. Here, we develop a model describing the population dynamics of a conspicuous defended prey species, and we assess the invasion conditions of derived phenotypes that differ from the ancestral phenotype by their conspicuousness. Such change in conspicuousness may then modify their level of mimicry with the local community of defended species. Derived colour pattern displayed in this focal population can therefore be either exactly similar, partially resembling or completely dissimilar to the local mimicry ring displaying the ancestral colour pattern. We assume that predation risk depends (1) on the number of individuals sharing a given colour pattern within the population, (2) on the occurrence of co-mimetic defended species, and (3) on the availability of alternative edible prey. Using a combination of analytical derivations and numerical simulations, we show that less conspicuous colour patterns are generally favoured within mimicry rings, unless reduced conspicuousness impairs mimicry. By contrast, when a mutation affecting the colour pattern leads to a shift toward a better protected mimicry ring, crypsis is no longer necessarily beneficial and a more conspicuous colour pattern can be favoured. The selected aposematic pattern then depends on the local composition of mimetic communities, as well as on the detectability, memorability and level of mimicry of the colour patterns.

## INTRODUCTION

The evolution of colour patterns in defended species is puzzling because conspicuousness, which determines the detectability of prey by predators, associates with multiple costs (Ruxton, 2019). Notably, prey individuals may suffer from increased attack risk by predators stemming from higher detectability of their more conspicuous colour patterns (Mappes *et al.*, 2014; Arias *et al.*, 2019), even if those prey are defended (Srygley and Kingsolver, 1998). However, selection against conspicuous colourations can be counter-balanced by the increased resemblance of aposematic patterns to the local prey communities, which may favour the evolution of conspicuousness in defended species. Indeed, Müllerian mimicry, whereby different defended prey species living in sympatry share the same colour pattern, reduces the individual predation risk (Müller, 1879). After experiencing the defences of a prey belonging to a given mimicry ring, predators tend to avoid prey displaying the same colour pattern. The protection gained by prey with a different level of conspicuousness then depends on the level of similarity of the colour pattern they displayed to the local mimicry rings, and on the generalization behaviour of predators (Kikuchi & Pfennig, 2010; Merrill *et al.*, 2012; Chouteau *et al.*, 2016).

When a derived colour pattern with a different level of conspicuousness emerges in a prey population, it can be perceived by predators as partly or totally different from the ancestral colour pattern, thereby increasing predation risk (Greenwood *et al.*, 1989; Lindström *et al.*, 2001). By contrast, the evolution of conspicuousness of colour pattern within a species might be facilitated when it results in a shift to an alternative mimicry ring that increases protection against predators. Such shift happens when a mutation produces a colour pattern that no longer resembles the ancestral colour pattern, but rather matches the colour pattern displayed within an already existing alternative mimicry ring. Individuals with a different conspicuousness of colour pattern can therefore either (1) be perceived by predators as similar to the ancestral mimicry ring, (2) be considered by predators as more similar to an alternative mimicry ring or (3) perceived by predators as different from all local mimicry ring.

A change in conspicuousness may not necessarily modify the signal recognized by predators: some other features of the colour pattern can be efficient in triggering predator memorability, and therefore predator avoidance learning, whatever the level of conspicuousness of the colour pattern. Alternatively, conspicuousness can be the most memorable feature of the aposematic colour pattern, such that only highly conspicuous colour pattern triggers rapid avoidance learning in predators. Altogether, memorability, crypsis and mimicry can therefore shape the evolution of conspicuousness. The defended butterflies from the Ithomiini tribe is a striking example where those three components are likely to affect the evolution of conspicuousness. These butterflies exhibit mildly-conspicuous aposematic signals, where wings are composed of cryptic transparent parts, combined with a few coloured elements (Corral‐Lopez *et al.*, 2021). In those aposematic butterflies, transparency decreases detectability by avian predators (Arias *et al.*, 2019; McClure *et al.*, 2019). Yet, mimicry among Ithomiini species and with other lepidoptera suggests that those mildly conspicuous colour patterns are also under selection for convergence and therefore truly act as aposematic signals (Beccaloni, 1997). Those different selection pressures could explain the persistence of transparent cryptic wing pattern associated with some key memorable features observed in Ithomiini clearwing species. Overall, selection pressure on conspicuousness within mimicry rings may ultimately influence Müllerian mimetic interactions by favouring reduced detectability in some mimetic species, thereby diminishing predation risk on less noticeable mimetic prey (Arias *et al.*, 2019).

Traditionally, modelling studies investigating the evolution of aposematism do not consider mimetic interactions (Leimar *et al.*, 1986; Speed & Ruxton, 2005a, 2005b; Broom *et al.*, 2006), while those investigating the evolution and implications of mimicry generally consider all colour patterns to have the same impact on predator learning behaviour (Müller, 1879; Sherratt, 2006; Gompert *et al.*, 2011; Aubier *et al.*, 2017; but see Franks *et al.*, 2009). Our theoretical study is a first step to fill this gap. Here we investigate the evolution of conspicuousness, by focusing on the interplay between protection provided by co-mimetic communities, and specific properties of the colour pattern itself, such as detectability and memorability (see Fig. 1 for an illustration). We use a mathematical modelling approach to test whether a mutation affecting the conspicuousness of the colour pattern can invade in a defended species engaged in Müllerian mimicry, depending on the effect of the mutation on (1) the phenotypic similarity to different mimicry rings, and on (2) the detectability and memorability of the derived colour pattern. Because the availability of alternative prey has been shown to be an important factor influencing the evolution of mimicry (Kokko *et al.*, 2003), we also investigate how the presence of alternative edible prey influence the evolution of conspicuousness, alongside the presence of co-mimetic species.

**Figure 1:**
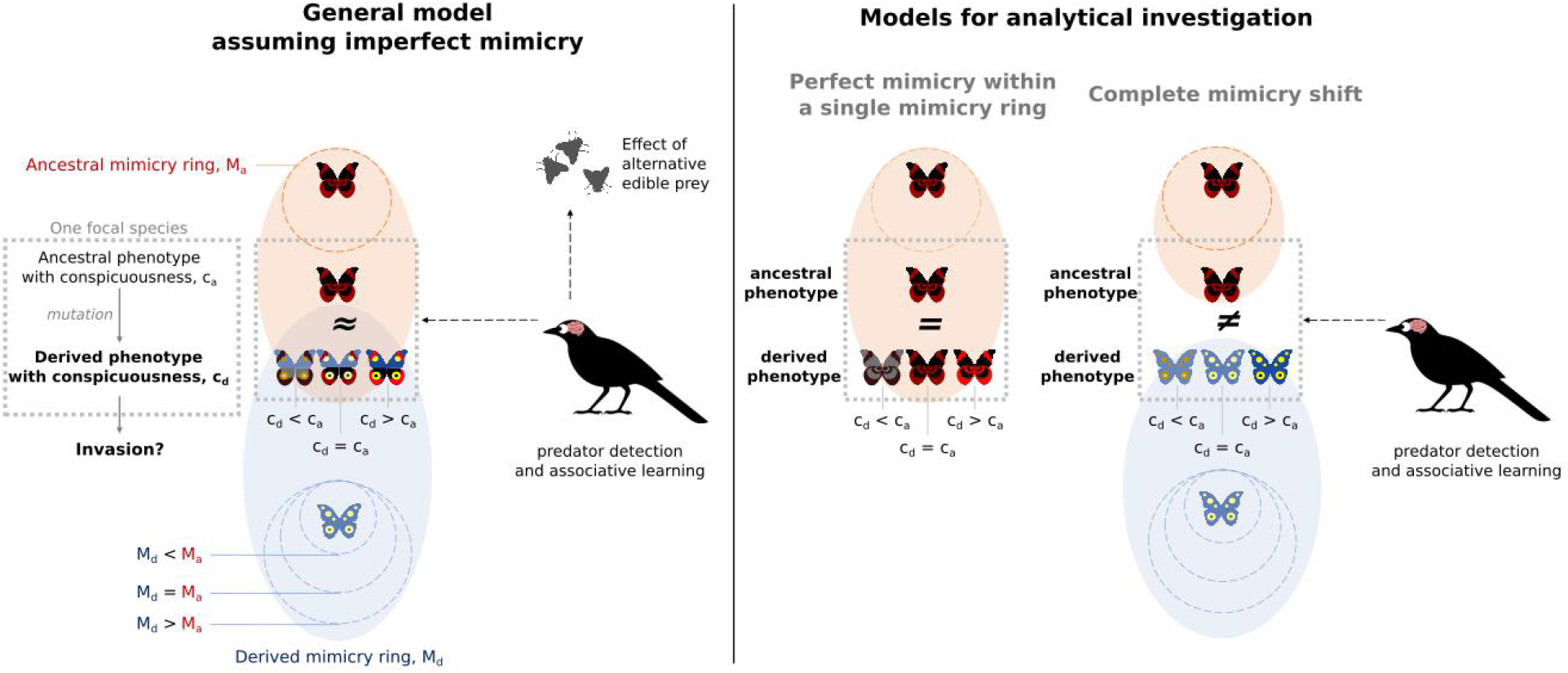
General scheme of our model investigating the evolutionary fate of more conspicuous colour pattern in a population of prey involved in Müllerian mimicry. We model the population dynamics of a single focal species composed of individuals displaying either the ancestral phenotype with conspicuousness c_a_ or the derived phenotype with conspicuousness c_d_. We test whether a mutation generating a derived phenotype with a different conspicuousness level than the ancestral phenotype can invade the population. Variation in conspicuousness can lead to different effect on the recognition of the colour patterns by the predator community. In our general model, we assume that the derived phenotype is only partially similar to the ancestral phenotypes and to the ‘ancestral mimicry ring’ displaying the ancestral colour pattern (imperfect mimicry), and we consider that alternative edible prey may alter the incentive of predators to attack cryptic prey. We also consider that the derived phenotype may match an alternative mimicry ring that we call ‘derived mimicry ring’. For analytical investigation, we consider two scenarios: 1) ‘perfect mimicry within a single mimicry ring’: both ancestral and derived phenotypes are accurate mimics to each other and to the local mimicry ring displaying the ancestral colour pattern, and 2) ‘complete mimicry shift’: the derived phenotype is totally dissimilar to the ancestral phenotypes and can be similar to an alternative community of defended species (simplified equations [9]). Note that our model thus explores all possible combinations of conspicuousness and mimicry levels in the ancestral and derived phenotypes. The population dynamic of predators is not explicitly modelled, but we do consider their perception and memory capacities.

## MATERIAL AND METHODS

### General model

Using ordinary differential equations, we model the population dynamics of a conspicuous defended prey species. This focal species is composed of individuals all harbouring the same level of defence and displaying a colour pattern phenotype (Fig. 1). Each individual can either display the *ancestral* phenotype (hereafter referred to using the subscript ‘*a*’) or the *derived* phenotype (referred to using the subscript ‘*d*’). Individuals with ancestral and derived phenotypes can differ in their conspicuousness level, *c*_*a*_ and *c*_*d*_. Variations in conspicuousness can affect the perception of the colour pattern by predators, so that the derived phenotype can be either perceived as (1) totally similar to the ancestral phenotype (perfect mimicry), (2) partially similar to the ancestral phenotype (imperfect mimicry) or (3) totally different from the ancestral phenotype. In these first two cases, the derived phenotype may also benefit from the protection provided by the presence of co-mimetic species matching the ancestral colour pattern (referred to as the ‘ancestral mimicry ring’). In these three cases, the derived phenotype may also benefit from the protection provided by an alternative mimicry ring that may look similar to the derived colour pattern (‘derived mimicry ring’; note that this terminology does not necessarily imply that this mimicry ring provides a lower protection than the ‘ancestral mimicry ring’).

Following Joron & Iwasa (2005), the changes through time in densities *N*_*a*_ and *N*_*d*_ of individuals displaying the ancestral and derived phenotypes depend on both local demography (described by the term *R*_*a*_ and *R*_*d*_) and predation (described by the terms *P*_*a*_ and *P*_*d*_):

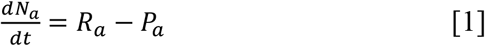

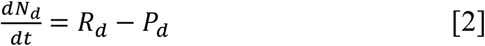

### Local demography

Individuals displaying the ancestral and derived phenotypes belong to the same species. As such, they share the same baseline growth rate *r*, compete for the same resources and share the same carrying capacity *K* (see Tab. 1 for a summary of the notations and default values). The changes in density of individuals displaying either the ancestral or derived colour pattern due to intra-specific competition follow a logistic regulation rule, as in the previous model of mimicry described by Joron & Iwasa (2005):

**Table 1.** Notations before rescaling the system of equations, and default values used in our model.

| Parameters and default values | Description |
| --- | --- |
| $N_a(t), N_d(t)$ | Number of individuals displaying the ancestral or derived phenotypes at time $t$ |
| $c_a, c_d$ | Conspicuousness of the ancestral or derived phenotypes |
| $M_a = 5,000$ | Reduction of the predation risk due to the ‘ancestral mimicry ring’ matching the ancestral phenotype (includes the abundance, defence, the conspicuousness and the memorability of the colour patterns carried by those species) |
| $M_d$ | Reduction of the predation risk due to the ‘alternative mimicry ring’ matching the derived phenotype (includes the abundance, defence, the conspicuousness and the memorability of the colour patterns carried by those species) |
| $\beta_a = \beta_d = 1$ | Translation parameters modulating the effect of conspicuousness on the memorability of the ancestral or derived phenotypes by predator |
| $u = 1$ | Associative learning response by predators due to the defence levels of individuals in the focal population |
| $p = 0.5$ | Baseline predation rate |
| $h = 0$ | Availability of alternative edible prey modulating the predation risk applied to by cryptic phenotype in the focal defended population (with $c_a$ or $c_d$ equal to 0) |
| $l = 0$ | Phenotypic distance between the ancestral and derived phenotypes that is not related to differences in conspicuousness |
| $\gamma$ | Perceived dissimilarity between ancestral and derived phenotypes, after accounting for the generalization behaviour of predators |
| $S$ | Level of similarity between ancestral or derived phenotypes; function of $\gamma$, $l$, $c_a$ and $c_d$ |
| $r = 10$ | Baseline growth rate of the focal species |
| $K = 1,000$ | Carrying capacity of the focal species |

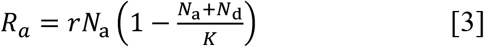

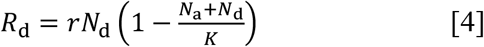

### Predation

Individuals with ancestral or derived phenotypes are characterized by their conspicuousness *c*_*a*_ and *c*_*d*_, respectively. Their predation risk depend on their colour pattern phenotype and its associated characteristics (detectability and memorability), and on the composition of the local community of defended and edible prey. Specifically, we assume that all individuals from the focal species, whatever their phenotype, have the same level of defence, which triggers the same associative learning response in predators (through parameter *u*). The baseline predation rate, *p*, on individuals with alternative and derived phenotype is also modulated by their detectability (modulated by their conspicuousness *c*_*a*_ and *c*_*d*_), by their memorability (through parameters *β*_*a*_ and *β*_*d*_), and by the abundance of alternative edible prey (through parameter *h*) and the presence of local mimicry rings matching the ancestral and derived coloration (through parameters *M*_*a*_ and *M*_*d*_). The protection gained by individuals displaying a given phenotype is then modulated by the similarity, *S*, to the other individuals from the focal species (but also by the similarity to the local mimicry rings).

The predation rate for individuals displaying either the ancestral or derived phenotype is thus modelled as:

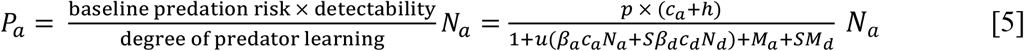

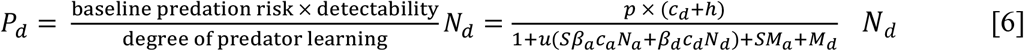

Importantly, we do not model explicitly the density of predators. Instead, we consider predation through those mortality functions, which capture how predators affect the mortality of individuals belonging to a focal species (Joron & Iwasa, 2005). The important features of these mortality functions are described below (see also Appendix A for more details).

#### Density-dependent predation risk

The predation risk is modelled following Joron & Iwasa (2005) and thus describes the density-dependent mortality risk incurred by any defended prey facing a natural predator community. Within this predator community, some predator individuals have already encountered the warning colour pattern and have learned to avoid it, while some other predator individuals are completely naïve (e.g. migrating individuals or juveniles) or are still learning about the warning colour pattern. As a result, the predation risk decreases as the number of individuals sharing the same colour pattern increases (Müller, 1879). We thus assume that the degree of predator learning increases as the density of individuals sharing the same colour pattern increases in equations [5] and [6].

#### Effects of conspicuousness on the predation risk

We assume that the conspicuousness of the colour pattern has two opposite effects on predation risk. On the one hand, increased conspicuousness increases the risk of being detected by predators, and therefore of being attacked (numerator terms in equations [5] – [6]). On the other hand, increased conspicuousness makes the colour pattern easier to remember for predators, and therefore enhances protection brought by predator learning (Sherratt, 2002; Broom *et al.*, 2006) (in the denominator terms in equations [5] – [6]).

Depending on the type of colour pattern variation (e.g., variations in colour contrast or pattern) triggering variations in conspicuousness, the mutation affecting conspicuousness can change the memorability of the colour pattern. The effect of conspicuousness on predator learning is modulated by parameters *β*_*a*_ and *β*_*d*_, which describe the levels of memorability induced in predators by given levels of conspicuousness (in the ancestral and derived phenotypes, respectively). Individuals with ancestral and derived phenotypes can thus differ in their levels of memorability, expressed as *β*_*a*_ × *c*_*a*_ and *β*_*d*_ × *c*_*d*_ respectively (in the denominator terms in equations [5] – [6]). If *β*_*a*_ = 0 or *β*_*d*_ = 0, the phenotype is not memorized even when it is conspicuous, and predator never associate the colour pattern with defence (*i.e.,* this is not a warning colour pattern). We therefore focus on cases where *β*_*a*_ > 0 and *β*_*d*_ > 0, *i.e.*, when an increase in conspicuousness associates with an increase in memorability (thereby triggering more rapid avoidance learning in predators and reducing predation).

#### Mimetic environment and shifts in mimicry ring

We consider that our focal conspicuous defended species is not isolated but instead belongs to a mimicry ring, as often observed in nature (Mallet & Gilbert, 1995). To keep the model analytically tractable, we do not model explicitly the population dynamics of each species of the mimicry community, just like we did not model explicitly the predator community. Instead, we model the protection provided by the local mimicry rings, taking the form of a higher degree of predator learning of the colour patterns carried by the focal individuals (in the denominator terms in equations [5] – [6]). Those mimicry rings can harbour either the ancestral or the derived colour pattern, through parameters *M*_*a*_ and *M*_*d*_, respectively. Parameters *M*_*a*_ and *M*_*d*_ thus capture the protection provided by co-mimetic defended prey, and therefore account for the abundance, the level of defence, the conspicuousness and the memorability of the mean phenotype of the co-mimetic species. By changing the level of conspicuousness, mutations in the focal species can also change the level of similarity to the ancestral mimicry ring, but it can also trigger a shift in colour pattern, resulting in a greater similarity of the derived phenotype to a different mimicry ring (providing a protection through *M*_*d*_ > 0).

#### Imperfect mimicry

We assume that variation in the level of conspicuousness between the ancestral and derived phenotypes can modify the level of similarity perceived by predators, thereby modulating predator generalization (Kikuchi & Pfennig, 2013; Motyka *et al.*, 2020). Mutations can thus generate derived colour patterns that are perceived as slightly different from the ancestral phenotype by the predators (*i.e.* imperfect mimicry). We thus model the degree of similarity *S* ∈ [0,1] between the ancestral and the derived phenotypes, which affects the recognition by predators. For *S* = 1, ancestral and derived phenotypes resemble each other and are perceived by predators as having the same phenotype. By contrast, for *S* = 0, ancestral and derived phenotypes do not look alike and are perceived by predators as completely distinct. Following Ruxton *et al.* (2008), the similarity level is defined as a Gaussian generalization function:

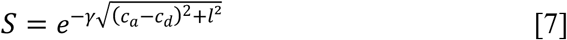

The parameter γ ∈ [0; +∞ [ describes the generalization behaviour of predators, *i.e.* how much they perceive phenotypic differences. The parameter *l* represents the distance between the ancestral and the derived colour patterns; *i.e.* the phenotypic distance that is not related to differences in conspicuousness. Hence, the phenotypic similarity between derived and ancestral phenotypes, as perceived by predators, depends on differences in colour pattern (*l*^2^) and differences in conspicuousness ( (*c*_*a*_ − *c*_*d*_)^2^). In our main analysis, however, we perform our analyses with *l* = 0 and we implement different values of *l* in supplementary analyses to investigate the effect of this parameter.

#### Availability of alternative prey

The availability of alternative edible prey may affect the effort of predators to seek for cryptic prey when resources are scarce. We thus introduce a parameter, *h*, which modulates the prey baseline mortality rate, so that even cryptic prey (with conspicuousness *c*_*a*_ or *c*_*d*_ equal to 0) can be attacked. The lack of alternative prey makes predators more motivated to search for cryptic prey, hence increasing the predation rate of cryptic prey. Therefore, the lack of alternative edible prey translates into high values of *h*. By contrast, a high abundance of alternative edible prey translates into low values of *h*.

### Analytical derivations under two scenarios: ‘perfect mimicry within a single mimicry ring’ and ‘complete mimicry shift’

We can derive analytical solutions for our system of equations, in two opposite scenarios:

1. ‘*Perfect mimicry within a single mimicry ring’* – Ancestral and derived phenotypes differ in conspicuousness but are perceived by predators as similar colour patterns (similarity *S* = 1; e.g., obtained for complete predator generalization γ = 0). Thus, individuals with either ancestral or derived phenotypes belong to the same mimicry ring that provides the same protection *M*_a_ + *M*_*d*_. Under this condition, it makes sense to consider that *M*_*d*_ = 0 so that *M*_*a*_ alone reflects the protection provided by the ancestral mimicry ring (in the denominator term in equations [5] and [6]). Ancestral and derived phenotypes nevertheless still differ in their conspicuousness and, in most cases, also in their memorability (*c*_*a*_ × *β*_*a*_ ≠ *c*_*d*_ × *β*_*d*_). For analytical tractability, we assume here that *h* = 0.
2. *‘Complete mimicry shift’* – Ancestral and derived phenotypes differ in conspicuousness and derived phenotypes display a colour pattern perceived as completely different from the ancestral one by predators (similarity *S* = 0; e.g., obtained for the absence of predator generalization *γ* → ∞). The ancestral and derived phenotypes are thus not generalized by predators, but the derived phenotype may match a different mimicry ring characterized by parameter *M*_*d*_ (in the denominator term in equation [6]). Because ancestral and derived phenotypes are distinct colour patterns, they may also have a different associated memorability. For analytical tractability, we also assume here that *h* = 0.

To derive analytically the system of equations of these two situations, we rescale all variables and parameters to baseline growth rate and carrying capacity, as follows: 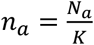, rescaled density of ancestral individuals; 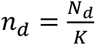, rescaled density of derived individuals; 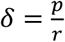, rescaled baseline mortality rate; *τ* = *rt* rescaled time unit; and *λ*_*a*_ = *uKβ*_*a*_ and *λ*_*d*_ = *uKβ*_*d*_ named rescaled deterrence factor (as in Joron & Iwasa 2005). Note that by defining the rescaled parameters *λ*_*a*_ and *λ*_*d*_, we combine the effects of unpalatability and memorization on the learning of the colour pattern by predators.

The dynamical system assuming the ‘Perfect mimicry within a single mimicry ring’ (*h* = 0, *M*_*d*_ = 0, and *S* = 1) then becomes:

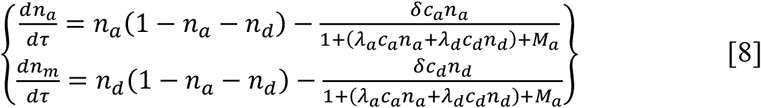

By contrast, assuming ‘Complete mimicry shift’ (*h* = 0 and *S* = 0), the system becomes:

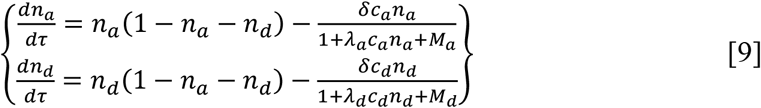

All analytical derivations are detailed in Appendix B. We investigate analytically the evolution of conspicuousness by conducting invasion analyses on the systems of equations [8] and [9]. We first derive the expression of the density 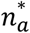 at equilibrium when the initial population is composed solely of ancestral individuals (solution of 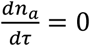, assuming *n*_*d*_ = 0). We then determine the sign of the growth rate of the derived population, assuming that derived phenotypes are rare within a population of individuals displaying the ancestral phenotype at equilibrium (sign of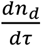, assuming 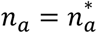 and *n*_*d*_ ≪ *n*_*a*_). If the sign of this growth rate is positive, then derived phenotypes can invade the population.

### Numerical analyses

We cannot get any analytical results from the general model, in particular when we account for imperfect mimicry (*i.e.* for intermediate value of phenotypic similarity *S* to local mimicry rings, 0 < *S* < 1; obtained for intermediate values of *γ*), or for increased predation rate on cryptic prey (for *h* > 0). In those cases, we assess the density 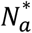 of the ancestral population at equilibrium numerically (reached for *t* = 1,000). We then infer invasion numerically by calculating the sign of the derivative 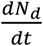 when 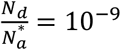 and by assuming that the derived phenotype can invade when 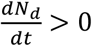.

Given that invading derived phenotypes will eventually replace the ancestral phenotypes in the population (as shown numerically in Supp. Fig. 1), we can infer the equilibrium (fixation of the ancestral or derived colour pattern) from the invasion analysis.

## RESULTS

### 1 A decrease in conspicuousness is favoured within a single mimicry ring (assuming ‘perfect mimicry within a single mimicry ring’; *S* = 1)

We first explore the evolution of conspicuousness when ancestral and derived phenotypes are perceived as perfectly similar by predators, and thus belong to the same mimicry ring (panels marked by a single symbol ‘*’ in Fig. 2). The derived phenotype only invades when its conspicuousness is lower than that of the ancestral phenotype: *c*_*d*_ < *c*_*a*_ (analytical derivations detailed in Appendix B).

**Figure 2:**
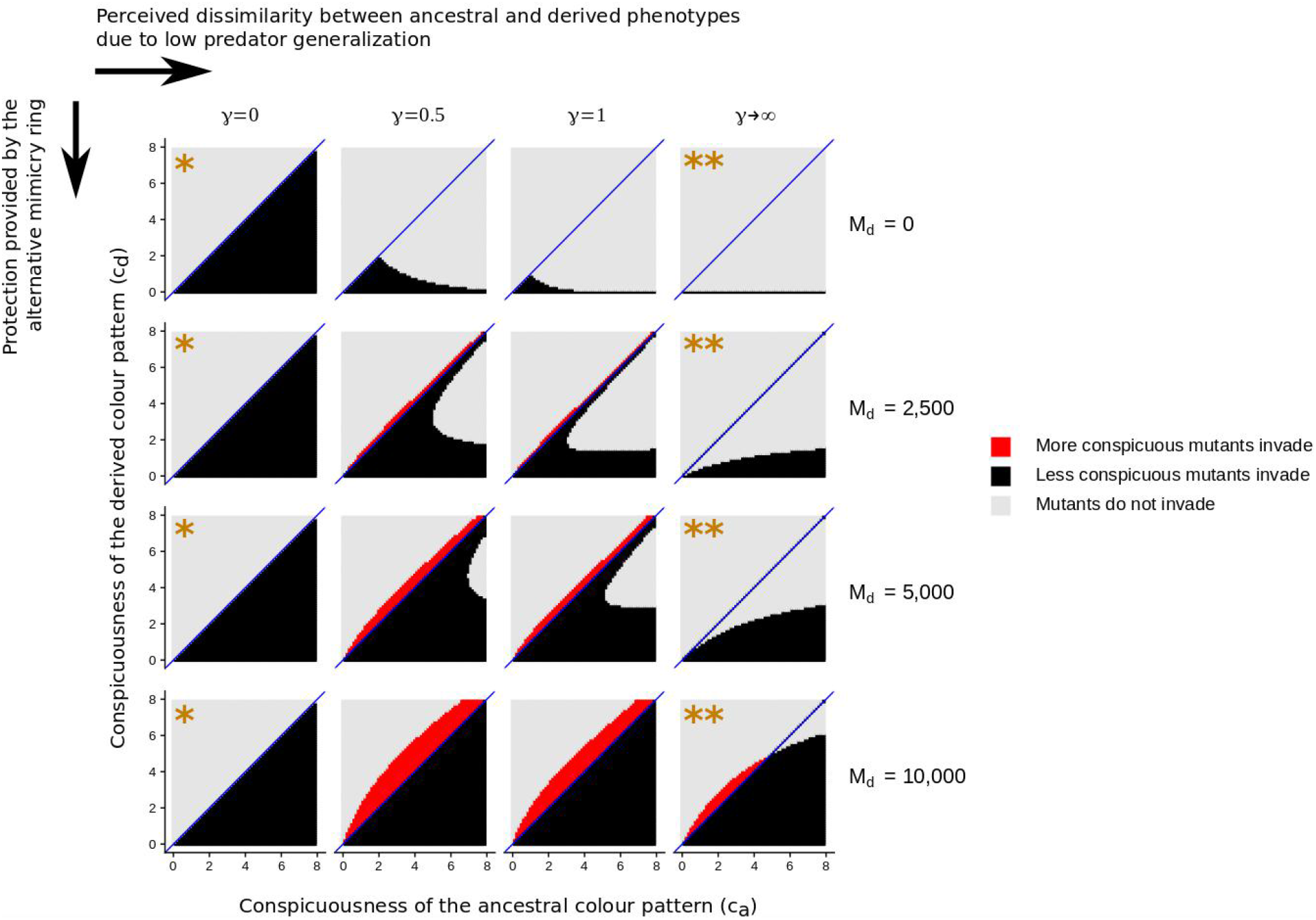
Evolution of conspicuousness depending on the mimetic community and on the similarity between ancestral and derived phenotypes. Inference of the evolutionary equilibrium from the invasion analysis, with ancestral and derived phenotypes having different levels of conspicuousness (*c*_*a*_, *c*_*d*_). We consider different values of parameter *M*_*d*_, which controls the protection provided by the alternative mimicry ring, and of parameter *γ*, which controls the perceived dissimilarity between ancestral and derived phenotypes (the condition *γ* → ∞ is obtained by assuming no similarity between phenotypes; i.e., S = 0). Note that the ancestral mimicry ring provides a protection *M*_*a*_ = 5,000. The diagonal blue line represents cases where *c*_*a*_ = *c*_*d*_. Colour scale indicates the sign of the derivative 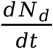 assuming mutants are rare. Light grey represents cases where the derived phenotype does not invade 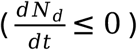, black represents cases where a less conspicuous phenotypes invades 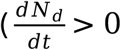 and *c*_*d*_ < *c*_*a*_ and red represents cases where a more conspicuous phenotypes invades 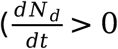 and *c*_*d*_ < *c*_*a*_). Symbols * and ** represent situations where we assessed analytically the conditions of invasions of a derived phenotype (‘perfect mimicry within a single mimicry ring’, and ‘complete mimicry shift’, respectively). When invasion occurs, it ultimately leads to the fixation of the derived phenotype (as shown in Supp. Fig. 1 with the same parameter combinations). Note that the panels on the left are all identical because protection *M*_*d*_ benefits to all individuals when mimicry is perfect (*γ* = 0); an increase in *M*_*d*_ therefore reflects an increase in *M*_*a*_. Here, *β*_*a*_ = 1 and *h* = 0. See other default values in Table 1.

Assuming perfect mimicry to the ancestral mimicry ring, selection therefore favours less conspicuous colour patterns. Such evolutionary process may occur via small-effect mutations modifying conspicuousness. Less conspicuous individuals are indeed less detectable while simultaneously benefiting from the protection provided by the mimicry ring, and therefore suffer less predation overall than more conspicuous individuals. Such positive selection on more cryptic derived phenotypes is likely to ultimately lead to highly cryptic colouration as the evolutionary stable strategy.

### 2 A mimicry shift can promote an increase in conspicuousness (assuming a ‘complete mimicry shift’; *S* = 0)

We then explore the evolution of conspicuousness when the derived colour pattern is perceived by predators as completely distinct from the ancestral colour pattern, and perfectly matches an alternative mimicry ring (panels marked by the symbol ‘**’ in Fig. 2). This is likely to apply when the mutation leads to drastic changes in colour pattern (via a large-effect mutation) and a mimicry ring shift.

Our analytical results detailed in Appendix B show that the level of conspicuousness under which the derived phenotypes are advantaged when introduced in the population mostly composed of individuals with the ancestral phenotype is:

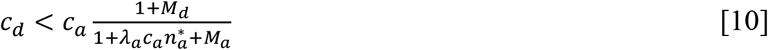

There is thus a conspicuousness threshold value under which the derived phenotypes are advantaged. The effects of parameters on this threshold value are summarized in Fig. 3 (see also Supp Tab. 1 and Supp. Fig. 2), and analytical derivations are detailed in Appendix B.

**Figure 3:**
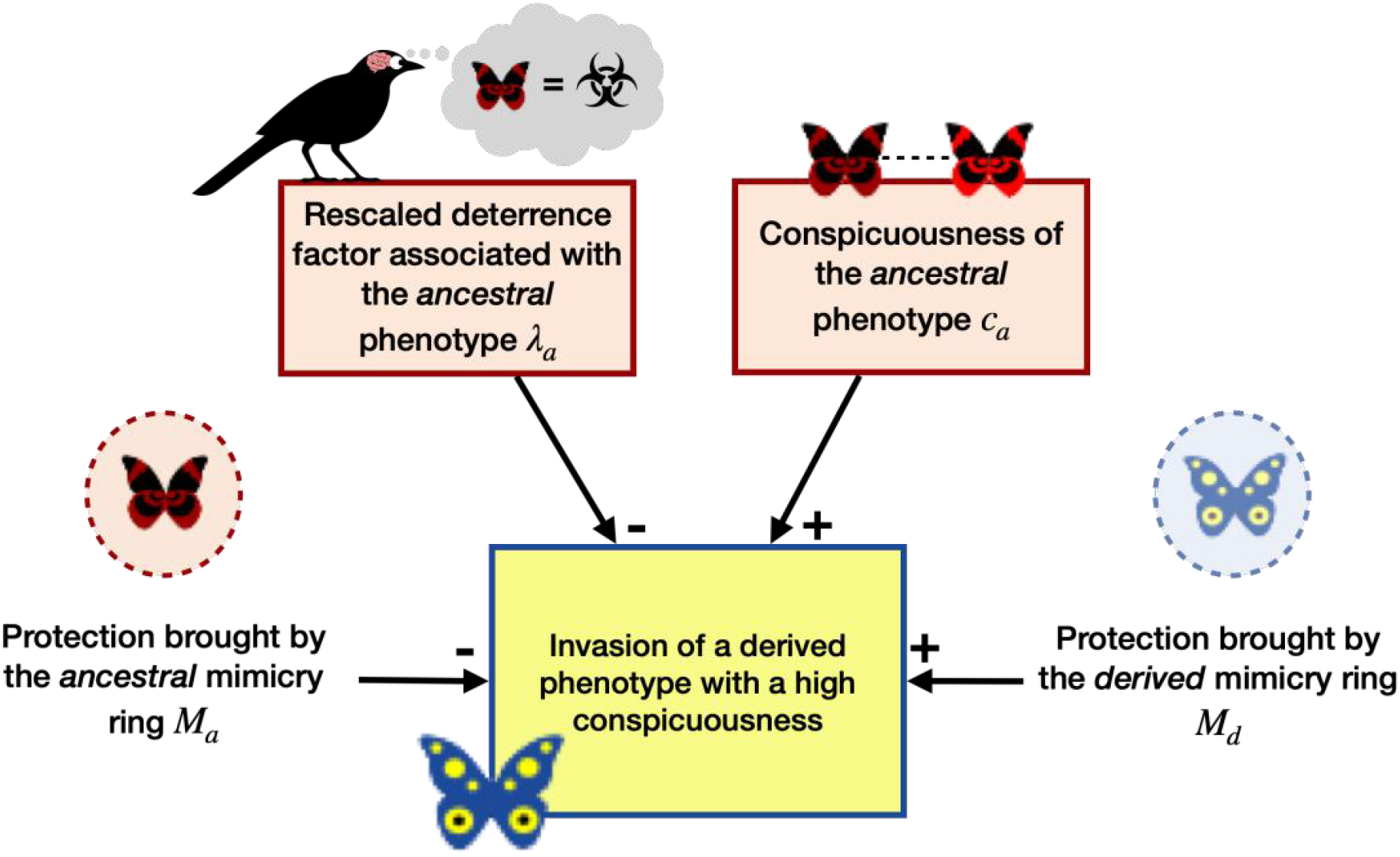
Effects of the parameters on the invasion of derived phenotypes, assuming a ‘complete mimicry shift’. Those effects are based on the effects of those parameters on the conspicuousness threshold 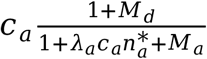 (see Eq. [10]). See also Supp. Tab. 1 and Supp. Fig. 2. Note that the rescaled deterrence factor associated with the ancestral phenotype is expressed as *λ*_*a*_ = *uKβ*_*a*_. The rescaled baseline mortality rate *δ* = *p/r* has a very little effect on the invasion of the derived phenotype and is therefore not represented here (Supp. Figs. 2 and 6).

Contrary to the results obtained above assuming ‘perfect mimicry within a single mimicry ring’, slightly less conspicuous derived phenotypes do not always invade, because of the potential costs associated with displaying a less protected alternative colour pattern (panels marked by a symbol ‘**’ in Fig. 2).

Overall, less conspicuous derived phenotypes are more advantaged than more conspicuous derived phenotypes. In particular, derived phenotypes that are completely cryptic (with *c*_*d*_ = 0; because we assume here that *h* = 0) always invades (because the threshold value 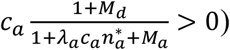), even when the derived colour pattern does not match any mimicry ring. Yet, derived phenotypes that are more conspicuous than the ancestral phenotype (*c*_*d*_ > *c*_*a*_) may be advantaged if the ratio shown in Eq. [10] is superior to 1, i.e., if 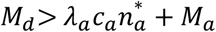 (area highlighted in red in the panels marked by a symbol ‘**’ in Fig. 2; see also Supp. Figs. 3 and 4). Consistent with our working hypothesis, our model confirms that for a more conspicuous derived colour pattern to be positively selected, it requires that the alternative mimicry ring (*M*_*d*_) provides better protection than the combination of the ancestral population 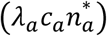 and the other species belonging to the same mimicry ring (*M*_*a*_). By contrast, when the mimicry ring matching the ancestral colour pattern provides a greater protection to ancestral individuals than does the other mimicry ring to the derived phenotype, this prevents derived colour patterns from invading (Fig. 3; see also Supp. Tab. S1 and Supp. Fig. 2).

### 3 Imperfect mimicry inhibits the invasion of derived phenotypes, and is more favourable to an increase in conspicuousness

We now investigate the effect of imperfect mimicry, by focusing on intermediate similarity between ancestral and derived phenotypes; with *S* ∈]0,1[ depending on the term *c*_*a*_ − *c*_*d*_; see Eq. [7]), assuming that perceived differences by predators only rely on conspicuousness (*l* = 0 and intermediate *γ*). Imperfect mimicry reduces not only the positive number-dependent selection brought by the ancestral individuals of the focal species on the derived phenotype, but also the protection from the entire ancestral mimicry ring on the derived phenotype. Imperfectly mimetic derived phenotypes are more favoured when their conspicuousness is similar to that of the ancestral phenotype, which increases their mimetic protection (*c*_*d*_ ≈ *c*_*a*_ for intermediate values of *γ* in Fig. 2, i.e., close to the blue diagonal; see also Supp. Figs. 3 and 4).

**Figure 4:**
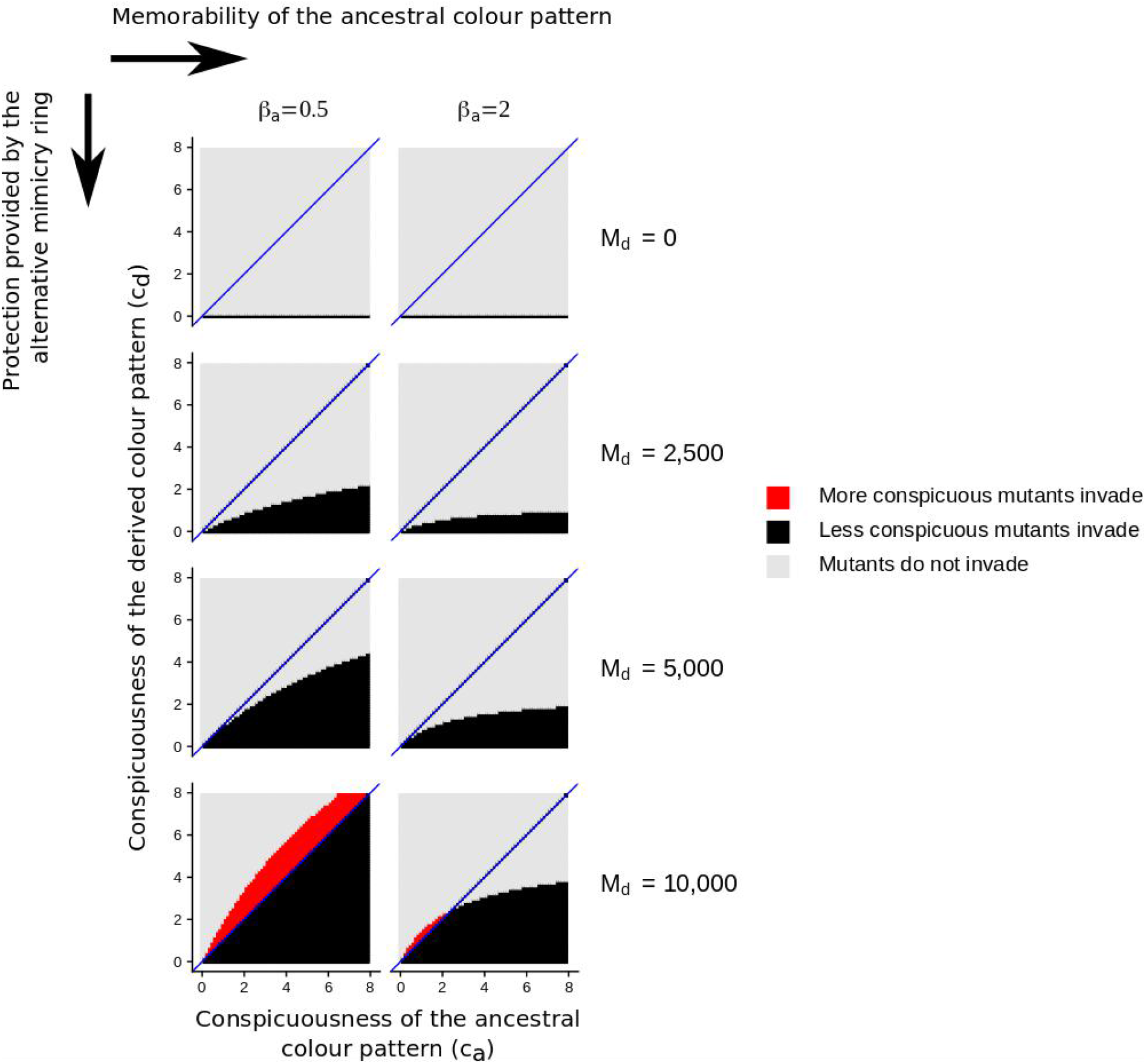
Evolution of conspicuousness depending on its impact on predator memorability. We consider different values of parameter *β*_*a*_, which controls the effect of conspicuousness on memorability in the ancestral phenotype. See Figure 2 for more details. Here, *γ* → ∞ (by setting *S* = 0) and *h* = 0. See other default values in Table 1.

When the imperfectly-mimetic derived phenotype matches an alternative mimetic community that provides strong protection (high *M*_*d*_; for intermediate values of *γ* in Fig. 2), the invasion of a more conspicuous derived colour pattern can occur easily when the derived conspicuousness remains similar to that of the ancestral phenotype (*c*_*d*_ ≈ *c*_*a*_). Imperfectly-mimetic derived phenotypes indeed benefit from resembling the ancestral phenotype while simultaneously roughly resembling another mimetic community, both reducing predation pressure. The fitness advantage associated with these *jack-of-all-trade* phenotypes (Sherratt 2002b) is confirmed in simulations for variable values of *l*, capturing the phenotypic distances that do not rely on conspicuousness and that are perceived by predators (i.e., differences in colour pattern; Supp. Figs. 3 and 5).

### 4 Easily-memorable phenotypes that facilitate predator learning inhibit the evolution of increased conspicuousness

The invasion of a more conspicuous derived phenotype when an alternative mimetic community co-occur is then modulated by the level of memorability induced by the ancestral colour pattern displayed by the focal species. We notice that when conspicuousness strongly enhances the memorability of the colour pattern (high values of *β*_*a*_; **Fig. 4 and Supp. Fig. 3**), the invasion of derived colour pattern is impaired just like when the defence triggers strong predator learning (high *u*) (Fig. 3 and Supp. Tab. S1). These two parameters indeed tune the intensity of the number-dependent selection driven by predator avoidance-learning (strength in number; *λ*_*a*_ = *uKβ*_*a*_), increasing the protection incurred to the ancestral phenotype. Contrastingly, the memorability associated with the derived colour pattern (controlled by *β*_*d*_; note however that parameter *M*_*d*_ also accounts for the memorability in co-mimetic species) does not affect the invasion conditions of the derived colour pattern, because derived phenotypes are rare initially and therefore *β*_*d*_*N*_*d*_ ≈ 0 (Supp. Fig. 4).

### 5 Predation pressure and availability of alternative prey modulate selection on conspicuousness

Increased baseline predation pressure (rescaled δ; Supp. Tab. S1) increases the range of conspicuousness values enabling the invasion of the derived phenotype: when predation rate is high, a slight decrease in conspicuousness in the derived phenotype is favoured, despite the cost associated with the shift to a less protected mimicry ring. Nonetheless, the magnitude of this effect is very low (Supp. Figs. 5 and 6). Therefore, a change in the abundance of alternative prey affects only weakly the evolution of conspicuousness, through a change in the baseline predation pressure.

The model also assumed that when alternative edible prey are scarce, predators have more incentive to search for cryptic prey (high values of *h*). In such condition, increased conspicuousness in the derived phenotype is more strongly favoured, a phenomenon enhanced by the protection provided by the derived mimicry ring (higher values of *M*_*d*_, Fig. 5; but note that the conspicuous values where higher conspicuousness is favoured does not change, as highlighted in Supp. Fig. 4). Our model thus highlights that increased conspicuousness in warning patterns is likely to be favoured when the baseline predation rate on cryptic prey is high, as expected when alternative food sources for predators are scarce.

**Figure 5.**
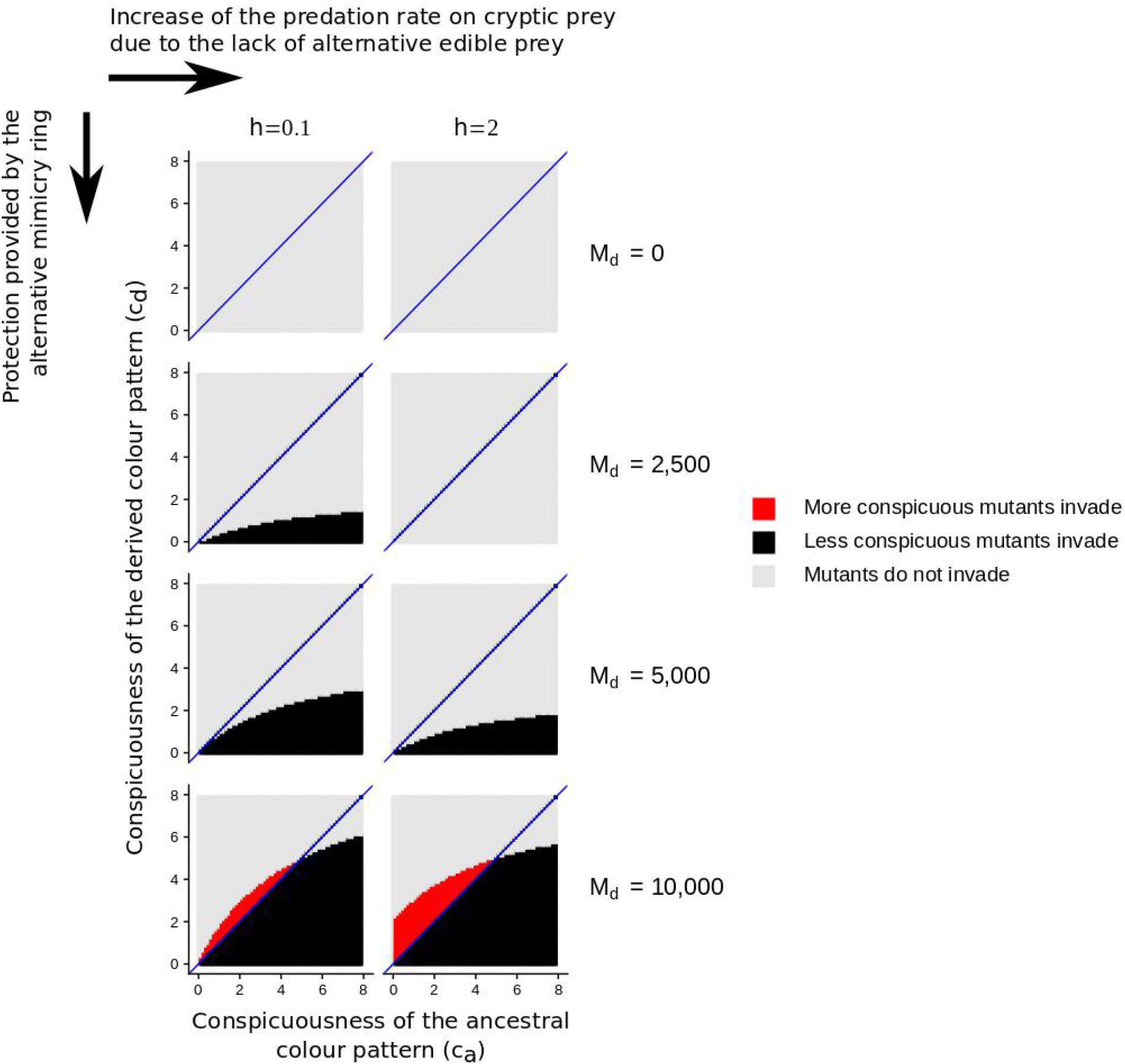
Evolution of conspicuousness depending on the abundance of alternative edible prey. We consider different values of parameter h, which controls the increase of the baseline predation rate on cryptic prey. See Figure 2 for more details. Here, *γ* → ∞ (by setting *S* = 0) and *γ*_*a*_ = 1. See other default values in Table 1.

## DISCUSSION

### Evolution toward decreasing level of conspicuousness within mimicry ring

Our model predicts that mimetic colour pattern may evolve toward reduced conspicuousness and complete crypsis within mimicry rings when mimicry is perfect (assuming ‘perfect mimicry within a single mimicry ring’ in our model). Less conspicuous individuals benefit from reduced detection by predators and therefore benefit from a lower predation rate than more conspicuous individuals. As long as the colour pattern is generalized by predators, all individuals within a mimicry ring benefit from the same reduction in predation risk, brought by the strength in number. Less conspicuous derived phenotypes act as selfish elements that may eventually replace the more conspicuous colour pattern, resulting in an increased predation risk for all members of the mimicry ring, because predator learning becomes less efficient as the mean conspicuousness decreases in the prey population. Ultimately, this would lead to completely cryptic colour pattern that is no longer detected by predators. This result is consistent with the higher rates of shifts from aposematism to crypsis than the reverse along the phylogeny of amphibians (Arbuckle & Speed 2015).

Nevertheless, variations of conspicuousness in natural colour pattern may result in colour pattern discrimination by predators, which limits the evolution of reduced conspicuousness within mimicry rings (a feature explored in our model when we assume imperfect mimicry; for intermediate *γ* and *M*_*d*_ = 0 a derived phenotype that is less conspicuous than the ancestral phenotype does not necessarily invade in Fig. 2). Such colour pattern discrimination could explain the persistence of key memorable conspicuous elements associated with transparent cryptic wings observed in Ithomiini clearwing species and their co-mimics (Beccaloni, 1997). Overall, selection pressure within mimicry rings may ultimately influence Müllerian mimetic interactions by favouring reduced detectability in some mimetic species, thereby diminishing predation risk on less noticeable mimetic prey (Arias *et al.*, 2019).

### Evolution of more conspicuous colour patterns can be promoted by mimicry shifts

Our model also highlights that a shift to more conspicuous colour patterns can be favoured when derived colour pattern matches a better-protected mimicry ring, via both perfect and imperfect mimicry. By considering the Müllerian mimetic environment in our model, we show that the conspicuousness of a colour pattern does not evolve independently from other aposematic species co-occurring in sympatry. Indeed, the evolution of conspicuousness depends on the protection provided by local defended mimetic communities, and on the presence of local edible prey communities. Here, we considered that variations in conspicuousness can result in different combinations of effects on memorability and generalization of the colour pattern. Even when assuming that conspicuousness does not associate with a direct benefit (*i.e.* we considered a worst-case scenario for conspicuousness invasion where crypsis is favoured), a shift to a distinct mimicry ring can still promote increased conspicuousness. Thus, our model shows that the evolution of increased conspicuousness can be promoted by natural selection exerted by predators and this prediction is in line with a previous empirical finding. For instance, the colouration of the mildly-defended viceroy butterfly, *Limenitis archippus*, is thought to have evolved from a non-aposematic ancestral pattern to an aposematic one matching the colour pattern of *Danaus* species (Platt *et al.*, 1971; Prudic *et al.*, 2002; Mullen, 2006; Prudic & Oliver, 2008). The evolution of a more conspicuous pattern could thus stem from mimicry towards the well-defended Monarch butterflies living in sympatry.

In addition to mimicry, our model highlights that the evolution of conspicuousness depends on the availability of alternative prey, assuming that large amounts of palatable prey reduces the selection on aposematic signals, following Kokko *et al.* (2003). Models exploring the foraging behaviour of predators nevertheless suggest that their discrimination capacities depend on a more complex interplay between the profitability and colour pattern variation found in prey communities (Getty 1985). Several cognitive biases in predators, not modelled in our study, might also favour the invasion of more conspicuous warning colour patterns in defended species: biased predation (Kikuchi *et al.*, 2020), innate memorability, direct deterrence, diet conservatism (Marples *et al.*, 2005) or neophobia (Marples & Kelly, 1999; Aubier & Sherratt, 2015) have indeed been shown to promote the emergence of new warning patterns. Our study thus calls for attention to the effect of ecological interactions with other defended species, edible alternative prey, and of the diversity of colour pattern in sympatric species on the evolution of aposematism in defended species.

### The effect of mimetic interaction on memorability can contribute to the evolution of mimetic colour pattern

The positive number-dependent protection gained by aposematic individuals can be modulated by the memorability of the colour patterns, a feature that we included in our model. While our study focuses on the evolution of conspicuousness, variations in the effect of conspicuousness on predator memorability may have other important consequences for the evolution of colour patterns in mimetic communities. For instance, the high diversity of aposematic colour patterns observed within localities (Beccaloni, 1997; Joron & Mallet, 1998; Willmott *et al.*, 2017; Briolat *et al.*, 2018) could be maintained by trade-offs between defence, memorability of the colour pattern and abundances in the different mimicry rings. In our study, we considered mimetic interactions without specifically modelling the abundance, the level of defences, and the memorability of the aposematic signals of co-mimetic species. Nonetheless, our model constitutes a fundamental basis for such future theoretical research explicitly investigating the evolution of colour pattern in multiple co-mimetic species.

## CONCLUSION

Overall, our study brings theoretical evidence that the trade-off between detectability and memorability induced by the conspicuousness of a warning colouration influences the evolution of mimetic colour patterns in defended species. Our model shows that in a worst-case scenario, where conspicuousness does not bring any direct benefit (such as immediate deterrence), the evolution of colour pattern with increased conspicuousness can only occur when alternative edible prey are rare and when a shift to a distinct and better protected Müllerian mimicry rings occurs. Considering mimetic community therefore opens a new angle to explain the evolution of the wide diversity of colour patterns: via number-dependent selection at the community level, more conspicuous colour pattern can be favoured. Importantly, however, such mechanism at the origin of the evolution of higher conspicuousness in a species relies on the existence of conspicuousness in other species; other processes must come into play for aposematic signals to emerge in the first place. Further researches accounting for mimetic interactions and variations in memorability between warning colour pattern are needed to better understand the evolution of aposematism at the level of the prey community.

## ACKNOWLEDGEMENTS

We thank three anonymous reviewers for their comments that helped us improve our manuscript considerably.

## DATA ACCESSIBILITY

The model is implemented in C++ and the code will be made available from a Dryad Digital Repository.

## AUTHOR’S CONTRIBUTIONS

XX, XX, XX and XX designed the study. XX and XX performed preliminary analyses not presented in the paper. XX and XX wrote the mathematical model and solved it analytically. XX performed all additional simulations. XX, XX and XX wrote the manuscript with contributions from other co-authors. All authors participated in constructive discussions and approved manuscript final version.

