## Appendix A for "Uncovering the effects of Müllerian mimicry on the evolution of conspicuousness in colour patterns"

### Appendix A – Mortality due to predation

#### 0 Purpose

Here, we describe how the mortality caused by predation was implemented in the model, using a step-by-step explanation.

#### 1 Original equation (Joron & Iwasa, 2005)

When facing aposematic prey, predators need to sample a certain number of a prey with a given phenotype per unit time to learn and avoid this particular phenotype. As a result, the predation rate becomes a strongly nonlinear, hyperbolically decreasing function of relative prey density in the population. Under such nonlinear density-dependence, selection against very rare phenotypes should be enhanced at very low frequencies, and should weaken as frequency increases.

Following Joron & Iwasa (*Journal of Theoretical Biology* 2005, **237**:87-103; doi: 10.1016/j.jtbi.2005.04.005), we account for such nonlinear density-dependence and we consider that the baseline mortality factors $d$ is multiplied by $1/(1 + s\,N)$ which produces a hyperbolic decrease in mortality with the density $N$ of the focal phenotype when $s > 0$. Parameter $s$ therefore encompasses any characteristic that determines the efficiency of associative learning (level of toxins, and memorability of the signal).

The predation rate on individuals with the focal phenotype, is:

$$P = \frac{d\,N}{1 + s\,N} \tag{A1}$$

The numerator of this equation represents the baseline mortality rate due to predation, and the denominator represents the reduction in predation due to associative learning by predators (which is more efficient as the density of individuals with the focal phenotype increases).

#### 2 Perfect mimicry between ancestral and derived phenotypes

From equation 1, we can express the mortality of individuals (with ancestral or derived phenotypes) that are perfect mimics as follows:

$$P_{\mathrm{a}} = \frac{d\,N_{\mathrm{a}}}{1 + s\,(N_{\mathrm{a}} + N_{\mathrm{d}})} \tag{A2}$$

$$P_{\mathrm{d}} = \frac{d\,N_{\mathrm{d}}}{1 + s\,(N_{\mathrm{a}} + N_{\mathrm{d}})} \tag{A3}$$

If the predators cannot perceive a difference between the two phenotypes, then all individuals suffer the same predation risk per capita ($\frac{P_{\mathrm{a}}}{N_{\mathrm{a}}} = \frac{P_{\mathrm{d}}}{N_{\mathrm{d}}}$) and benefit from a lower predation risk due to mimicry (term $N_{\mathrm{a}} + N_{\mathrm{d}}$ in the denominator).

#### 3 Imperfect mimicry between ancestral and derived phenotypes

We can now express the mortality of individuals (with ancestral or derived phenotypes) that are imperfect mimics, with resemblance captured by a quantity $S$, as follow:

$$P_{\mathrm{a}} = \frac{d\,N_{\mathrm{a}}}{1 + s\,(N_{\mathrm{a}} + S\,N_{\mathrm{d}})} \tag{A4}$$

$$P_{\mathrm{d}} = \frac{d\,N_{\mathrm{d}}}{1 + s\,(S\,N_{\mathrm{a}} + N_{\mathrm{d}})} \tag{A5}$$

If $S = 0$, then ancestral and derived phenotypes are perceived as completely different by predators. If $S = 1$, then ancestral and derived phenotypes are perfect mimics.

We assume that the similarity level depends on the difference in conspicuousness, $c_a - c_d$, and is expressed as a Gaussian generalization function, as defined in the main text:

$$S = \exp\left(-\gamma\sqrt{(c_a - c_d)^2 - l^2}\right) \tag{A6}$$

The parameter $\gamma$ describes the generalization behaviour of predators, i.e. how much they perceive phenotypic differences. The parameter $l$ represents the distance between the ancestral and the derived colour patterns; i.e. the phenotypic distance that is not related to differences in conspicuousness. See Ruxton *et al.* (2008) (*Evolution*, **62**(11), 2913-2921) for more details on this function.

### 4 Mimicry rings

To assume that other comimetic species reduce the predation risk by enhancing predator learning, we include additional terms in the denominator. Remember that the denominator represents the reduction in predation due to associative learning by predators.

$$P_a = \frac{d}{1 + s\ (N_a + S\,N_d) + M_a + S\,M_d} \tag{A7}$$

$$P_a = \frac{d}{1 + s\ (S\,N_a + N_d) + S\,M_a + M_d} \tag{A8}$$

High parameters values of $M_a$ and $M_d$ lead to decreased predation risks. Those parameters therefore describes the efficiency of the mimicry ring (i.e., the comimetic defended species) in reducing predation risk by enhancing associative learning by predators. The accuracy of mimicry with those other defended species depends on the resemblance between the phenotype carried by individuals and the mean phenotypes displayed in the comimetic species. We thus assume that the contribution of $M_a$ and $M_d$ in reducing predation risks is modulated by a factor that determines this resemblance. For simplicity, we assume that a derived phenotype that is an imperfect mimic of the ancestral phenotype is an equally imperfect mimic of the co-mimetic individuals that resemble the ancestral phenotype. We make the same assumption to describe the resemblance between ancestral phenotype and individuals from the mimicry rings matching the derived phenotype.

### 5 Increased detectability due to high conspicuousness

We assume that high conspicuousness results in a high risk of being detected by predators, thereby increasing the baseline mortality factor. Therefore, individuals with ancestral and derived phenotypes may have different baseline mortality factors, $d_a$ and $d_d$. In particular we assume increased conspicuousness increases the baseline mortality factor:

$$d_a = p\,c_a \tag{A9}$$

$$d_d = p\,c_d \tag{A10}$$

$$P_a = \frac{d_a\,N_a}{1 + s\ (N_a + S\,N_d) + M_a + S\,M_d} \tag{A11}$$

$$P_d = \frac{d_d\,N_d}{1 + s\ (S\,N_a + N_d) + S\,M_a + M_d} \tag{A12}$$

### 6 Increased associative learning due to high conspicuousness

We assume that high conspicuousness may result in more efficient associative learning by predators. Therefore, individuals with ancestral and derived phenotypes are associated with different density-dependence factors, $s_a$ and $s_d$. In particular, we assume that increased conspicuousness increases the density-dependence factor depending on how memorable the colour pattern is (modulated by parameter $\beta_a$ and $\beta_d$):

$$s_a = u\,\beta_a\,c_a \tag{A13}$$

$$s_d = u\,\beta_d\,c_d \tag{A14}$$

$$P_{\mathrm{a}} = \frac{d_{\mathrm{a}}\,N_{\mathrm{a}}}{1 + (s_{\mathrm{a}}\,N_{\mathrm{a}} + S\,s_{\mathrm{d}}\,N_{\mathrm{d}}) + M_{\mathrm{a}} + S\,M_{\mathrm{d}}} \tag{A15}$$

$$P_{\mathrm{d}} = \frac{d_{\mathrm{d}}\,N_{\mathrm{d}}}{1 + (S\,s_{\mathrm{a}}\,N_{\mathrm{a}} + s_{\mathrm{d}}\,N_{\mathrm{d}}) + S\,M_{\mathrm{a}} + M_{\mathrm{d}}} \tag{A16}$$

### 7 Implications of alternative edible prey

The presence of alternative edible prey can affect the population dynamics by making predators more or less hungry. We assume that a parameter $h$ adjusts the prey baseline mortality rate, so that even highly cryptic prey (with $c_{\mathrm{a}}$ or $c_{\mathrm{d}}$ equal to 0) can be attacked. The parameter $h$ can reflect the abundance of alternative edible prey. Indeed, hungry predators have more incentive to search for cryptic prey compared to well-fed predators, thus $h = 0$ occurs when there is a lot of alternative prey and high $h > 0$ occurs when there are few alternative edible prey. Parameter $h$ therefore modulate the baseline mortality rate:

$$d_{\mathrm{a}} = p\,(c_{\mathrm{a}} + h) \tag{A17}$$

$$d_{\mathrm{d}} = p\,(c_{\mathrm{d}} + h) \tag{A18}$$

### 8 Final system of equation

Overall, we get the system of equations presented in the manuscript:

$$d_{\mathrm{a}} = p\,(c_{\mathrm{a}} + h) \tag{A19}$$

$$d_{\mathrm{d}} = p\,(c_{\mathrm{d}} + h) \tag{A20}$$

$$s_{\mathrm{a}} = u\,\beta_{\mathrm{a}}\,c_{\mathrm{a}} \tag{A21}$$

$$s_{\mathrm{d}} = u\,\beta_{\mathrm{d}}\,c_{\mathrm{d}} \tag{A22}$$

$$P_{\mathrm{a}} = \frac{d_{\mathrm{a}}\,N_{\mathrm{a}}}{1 + (s_{\mathrm{a}}\,N_{\mathrm{a}} + S\,s_{\mathrm{d}}\,N_{\mathrm{d}}) + M_{\mathrm{a}} + S\,M_{\mathrm{d}}} \tag{A23}$$

$$P_{\mathrm{d}} = \frac{d_{\mathrm{d}}\,N_{\mathrm{d}}}{1 + (S\,s_{\mathrm{a}}\,N_{\mathrm{a}} + s_{\mathrm{d}}\,N_{\mathrm{d}}) + S\,M_{\mathrm{a}} + M_{\mathrm{d}}} \tag{A24}$$

$$S = \exp\left(-\gamma\sqrt{(c_{\mathrm{a}} - c_{\mathrm{d}})^2 - l^2}\right) \tag{A25}$$
