## Appendix B for "Uncovering the effects of Müllerian mimicry on the evolution of conspicuousness in colour patterns"

### Appendix B – Analytical derivations

#### 0 Purpose

We here derive analytically the conditions under which a derived phenotype can invade in the population, assuming either 'perfect mimicry within a single mimicry ring' or a 'complete mimicry shift', as defined in the main text.

Assuming 'perfect mimicry within a single mimicry ring', the derived phenotype differs from the ancestral phenotype by its conspicuousness only. Assuming a 'complete mimicry shift', the derived phenotype differ from the ancestral phenotype by its conspicuousness but also by its colour pattern (as assumed by the existence of an alternative mimicry ring).

We perform here local stability analyses. First, we study the dynamical system without the derived phenotype and we derive the density of individuals carrying the ancestral phenotype at equilibrium (Section 1). Second, we derive the conditions under which individuals with a derived phenotype have a positive growth rate, assuming that they are very rare initially, and that the density of individuals carrying the ancestral phenotype is at its equilibrium value (Section 2 assuming 'perfect mimicry within a single mimicry ring', Section 3 assuming a 'complete mimicry shift')

#### 1 Study of the dynamical system without the derived phenotype

Without individuals carrying the derived phenotype ($n_{\mathrm{d}} = 0$), the systems of equations are identical assuming 'perfect mimicry within a single mimicry ring' and assuming a 'complete mimicry shift':

$$\frac{\mathrm{d}n_{\mathrm{a}}}{\mathrm{d}\tau} = n_{\mathrm{a}}(1 - n_{\mathrm{a}}) - \frac{\delta c_{\mathrm{a}} n_{\mathrm{a}}}{1 + \lambda_{\mathrm{a}} c_{\mathrm{a}} n_{\mathrm{a}} + M_{\mathrm{a}}} \tag{B1}$$

Let $n_{\mathrm{a}}^*$ be the density of individuals carrying the ancestral phenotype at equilibrium, *i.e.* satisfying the condition $\frac{\mathrm{d}n_{\mathrm{a}}}{\mathrm{d}\tau} = 0$. Here, we are placed in conditions under which the population composed only of individuals with the ancestral phenotype does not get extinct, *i.e.* $n_{\mathrm{a}}^* > 0$. We have :

$$n_{\mathrm{a}}^*(1 - n_{\mathrm{a}}^*) - \frac{\delta c_{\mathrm{a}} n_{\mathrm{a}}^*}{1 + \lambda_{\mathrm{a}} c_{\mathrm{a}} n_{\mathrm{a}}^* + M_{\mathrm{a}}} = 0 \tag{B2}$$

then

$$n_{\mathrm{a}}^* = 1 - \frac{\delta c_{\mathrm{a}}}{1 + \lambda_{\mathrm{a}} c_{\mathrm{a}} n_{\mathrm{a}}^* + M_{\mathrm{a}}} \tag{B3}$$

We will further use these expressions to simplify analytical calculations.

$$n_{\mathrm{a}}^* = 1 - \frac{1 + \lambda_{\mathrm{a}} c_{\mathrm{a}} + M_{\mathrm{a}} - \sqrt{(1 + \lambda_{\mathrm{a}} c_{\mathrm{a}} + M_{\mathrm{a}})^2 - 4\lambda_{\mathrm{a}} c_{\mathrm{a}}^2 \delta}}{2\lambda_{\mathrm{a}} c_{\mathrm{a}}} \tag{B4}$$

$$n_{\mathrm{a}}^* = \frac{-(1 + M_{\mathrm{a}} - \lambda_{\mathrm{a}} c_{\mathrm{a}}) + \sqrt{(1 + M_{\mathrm{a}} - \lambda_{\mathrm{a}} c_{\mathrm{a}})^2 + 4\lambda_{\mathrm{a}} c_{\mathrm{a}}(1 + M_{\mathrm{a}} - \delta c_{\mathrm{a}})}}{2\lambda_{\mathrm{a}} c_{\mathrm{a}}} \tag{B5}$$

We are only interested in the case where $n_{\mathrm{a}}^* > 0$, which holds only when $1 + M_{\mathrm{a}} > \delta c_{\mathrm{a}}$.

***Proof:***

$n_{\mathrm{a}}^*$ exists, and $n_{\mathrm{a}}^* > 0$ if:

$$4\lambda_{\mathrm{a}} c_{\mathrm{a}}(1 + M_{\mathrm{a}} - \delta c_{\mathrm{a}}) > 0 \tag{B6}$$

Which is equivalent to:

$$1 + M_\mathrm{a} > \delta c_\mathrm{a} \tag{B7}$$

### 2 Invasion conditions assuming a 'complete mimicry shift'

We assume that the density of individuals carrying the derived phenotype is initially very low, *i.e.* we consider that $n_\mathrm{d} = O(\epsilon)$ with epsilon being small. To determine whether having the derived phenotype is advantageous or not, we determine the sign of the derivative of the density of individuals with the derived phenotype when $n_\mathrm{a} = n_\mathrm{a}^*$. We note $f(n_\mathrm{a}, n_\mathrm{d})$ the derivative of the density of individuals with the derived phenotype: $f(n_\mathrm{a}, n_\mathrm{d}) = \frac{\mathrm{d}n_\mathrm{d}}{\mathrm{d}\tau}$.

Hence, when $n_\mathrm{a} = n_\mathrm{a}^*$, we have:

$$f(n_\mathrm{a}^*, n_\mathrm{d}) = \frac{\mathrm{d}n_\mathrm{d}}{\mathrm{d}\tau}\Big|_{n_\mathrm{a}=n_\mathrm{a}^*} \tag{B8}$$

$$= n_\mathrm{d}(1 - n_\mathrm{a}^* - n_\mathrm{d}) - \frac{\delta c_\mathrm{d} n_\mathrm{d}}{1 + \lambda_\mathrm{a}(c_\mathrm{a} n_\mathrm{a}^* + c_\mathrm{d} n_\mathrm{d}) + M_\mathrm{a}} \tag{B9}$$

Given that $n_\mathrm{d} = O(\epsilon)$, we can approximate:

$$f(n_\mathrm{a}^*, n_\mathrm{d}) = n_\mathrm{d}(1 - n_\mathrm{a}^*) - \frac{\delta c_\mathrm{d} n_\mathrm{d}}{1 + \lambda_\mathrm{a} c_\mathrm{a} n_\mathrm{a}^* + M_\mathrm{a}} + O(\epsilon^2) \tag{B10}$$

By using equation B3, we get:

$$f(n_\mathrm{a}^*, n_\mathrm{d}) = \frac{\delta n_\mathrm{d}}{1 + \lambda_\mathrm{a} c_\mathrm{a} n_\mathrm{a}^* + M_\mathrm{a}}(c_\mathrm{a} - c_\mathrm{d}) + O(\epsilon^2) \tag{B11}$$

By neglecting the term of the same order as $\epsilon^2$, we find that having the derived phenotype is advantageous when:

$$c_\mathrm{d} < c_\mathrm{a} \tag{B12}$$

### 3 Invasion conditions assuming a 'complete mimicry shift'

Assuming a 'complete mimicry shift', when $n_\mathrm{a} = n_\mathrm{a}^*$, we have:

$$f(n_\mathrm{a}^*, n_\mathrm{d}) = n_\mathrm{d}(1 - n_\mathrm{a}^* - n_\mathrm{d}) - \frac{\delta c_\mathrm{d} n_\mathrm{d}}{1 + \lambda_\mathrm{a} c_\mathrm{d} n_\mathrm{d} + M_\mathrm{d}} \tag{B13}$$

$$= n_\mathrm{d}(1 - n_\mathrm{a}^*) - \frac{\delta c_\mathrm{d} n_\mathrm{d}}{1 + M_\mathrm{d}} + O(\epsilon^2) \tag{B14}$$

By using equation B3, we have :

$$f(n_\mathrm{a}^*, n_\mathrm{d}) = \delta n_\mathrm{d}\left(\frac{c_\mathrm{a}}{1 + \lambda_\mathrm{a} c_\mathrm{a} n_\mathrm{a}^* + M_\mathrm{a}} - \frac{c_\mathrm{d}}{1 + M_\mathrm{d}}\right) + O(\epsilon^2) \tag{B15}$$

By neglecting the term of the same order as $\epsilon^2$, we find that having the derived phenotype is advantageous when:

$$\frac{c_\mathrm{d}}{1 + M_\mathrm{d}} < \frac{c_\mathrm{a}}{1 + \lambda_\mathrm{a} c_\mathrm{a} n_\mathrm{a}^* + M_\mathrm{a}} \tag{B16}$$

and therefore:

$$c_\mathrm{d} < c_\mathrm{a}\frac{(1 + M_\mathrm{d})}{1 + \lambda_\mathrm{a} c_\mathrm{a} n_\mathrm{a}^* + M_\mathrm{a}} \tag{B17}$$

We call $\hat{C}$ this threshold value:

$$\hat{C} = c_\mathrm{a}\frac{(1 + M_\mathrm{d})}{1 + \lambda_\mathrm{a} c_\mathrm{a} n_\mathrm{a}^* + M_\mathrm{a}} \tag{B18}$$

##### 3.1 Sensitivity of $\hat{\mathbf{C}}$

The threshold value $\hat{C}$ below which the derived phenotype can invade can be simplified as:

$$\hat{C} = \frac{2\,(1 + M_\mathrm{d})\,c_\mathrm{a}}{1 + \lambda_\mathrm{a}\,c_\mathrm{a} + M_\mathrm{a} + \sqrt{X}} \tag{B19}$$

With:

$$X = (1 + M_\mathrm{a} + \lambda_\mathrm{a}\,c_\mathrm{a})^2 - 4\,\lambda_\mathrm{a}\,\delta\,{c_\mathrm{a}}^2 \tag{B20}$$

We now determine the sensitivity of this threshold value to a change in parameter values, to determine under what conditions a phenotype characterized by a high conspicuousness $c_\mathrm{d} > c_\mathrm{a}$ can invade (which occurs when the threshold value $\hat{C}$ is high). See Supp. Tab. S1.

**Effect of $\mathbf{M_d}$ on the threshold value $\hat{\mathbf{C}}$:**

$$\frac{\partial \hat{C}}{\partial M_\mathrm{d}} = \frac{2\,c_\mathrm{a}}{1 + \lambda_\mathrm{a}\,c_\mathrm{a} + M_\mathrm{a} + \sqrt{X}} > 0 \tag{B21}$$

Therefore, increased $M_\mathrm{d}$ increases the invasibility area. Interestingly, the derived phenotype with a higher conspicuousness than the ancestral phenotype can invade the population if:

$$M_\mathrm{d} > M_\mathrm{a} + \lambda_\mathrm{a} c_\mathrm{a} n_\mathrm{a}^*. \tag{B22}$$

**Effect of $\mathbf{M_a}$ on the threshold value $\hat{\mathbf{C}}$:**

$$\frac{\partial \sqrt{X}}{\partial M_\mathrm{a}} = \frac{1 + M_\mathrm{a} + \lambda_\mathrm{a}\,c_\mathrm{a}}{\sqrt{X}} > 0 \tag{B23}$$

$$\frac{\partial \hat{C}}{\partial M_\mathrm{a}} = \frac{-2(1 + M_\mathrm{d})c_\mathrm{a}}{\left(1 + \lambda_\mathrm{a}\,c_\mathrm{a} + M_\mathrm{a} + \sqrt{X}\right)^2}\left(1 + \frac{\partial \sqrt{X}}{\partial M_\mathrm{a}}\right) < 0 \tag{B24}$$

Therefore, increased $M_\mathrm{a}$ decreases the invasibility area.

**Effect of $\boldsymbol{\lambda_\mathrm{a}}$ on the threshold value $\hat{\mathbf{C}}$:**

$$\frac{\partial \sqrt{X}}{\partial \lambda_\mathrm{a}} = \frac{c_\mathrm{a}\,\left[1 + M_\mathrm{a} + (\lambda_\mathrm{a} - 2\,\delta)\,c_\mathrm{a}\right]}{\sqrt{X}} \tag{B25}$$

$$\frac{\partial \hat{C}}{\partial \lambda_\mathrm{a}} = \frac{-2\,(1 + M_\mathrm{d})\,{c_\mathrm{a}}^2}{\left(1 + \lambda_\mathrm{a}\,c_\mathrm{a} + M_\mathrm{a} + \sqrt{X}\right)^2\sqrt{X}}\left(\sqrt{X} + 1 + M_\mathrm{a} + \lambda_\mathrm{a}\,c_\mathrm{a} - 2\,\delta\,c_\mathrm{a}\right) \tag{B26}$$

$$\frac{\partial \hat{C}}{\partial \lambda_\mathrm{a}} = \frac{-2\,(1 + M_\mathrm{d})\,{c_\mathrm{a}}^2}{\left(1 + \lambda_\mathrm{a}\,c_\mathrm{a} + M_\mathrm{a} + \sqrt{X}\right)^2\sqrt{X}}\left(\sqrt{X} - (1 + M_\mathrm{a} - \lambda_\mathrm{a}\,c_\mathrm{a}) + 2\,(1 + M_\mathrm{a} - \delta\,c_\mathrm{a})\right) \tag{B27}$$

$$\frac{\partial \hat{C}}{\partial \lambda_\mathrm{a}} = \frac{-2\,(1 + M_\mathrm{d})\,{c_\mathrm{a}}^2}{\left(1 + \lambda_\mathrm{a}\,c_\mathrm{a} + M_\mathrm{a} + \sqrt{X}\right)^2\sqrt{X}}\left(2\lambda_\mathrm{a} c_\mathrm{a} n_\mathrm{a}^* + 2\,(1 + M_\mathrm{a} - \delta\,c_\mathrm{a})\right) \tag{B28}$$

Yet, $1 + M_\mathrm{a} - \delta\,c_\mathrm{a} > 0$ (imposed by the condition of existence of the equilibrium), and $\sqrt{X} - (1 + M_\mathrm{a} - \lambda_\mathrm{a}\,c_\mathrm{a}) > 0$. Therefore:

$$\frac{\partial \hat{C}}{\partial \lambda_\mathrm{a}} < 0 \tag{B29}$$

Therefore, increased $\lambda_\mathrm{a}$ decreases the invasibility area.

**Effect of $\delta$ on the threshold value $\hat{\mathbf{C}}$:**

$$\frac{\partial \sqrt{X}}{\partial \delta} = \frac{-2\,\lambda_\mathrm{a}\,{c_\mathrm{a}}^2}{\sqrt{X}} < 0 \tag{B30}$$

$$\frac{\partial \hat{C}}{\partial \delta} = \frac{-2\,(1+M_\mathrm{d})\,c_\mathrm{a}}{\left(1+\lambda_\mathrm{a}\,c_\mathrm{a}+M_\mathrm{a}+\sqrt{X}\right)^2}\,\frac{\partial \sqrt{X}}{\partial \delta} > 0 \tag{B31}$$

Therefore, increased $\delta$ increases the invasibility area.

**Effect of $\mathbf{c_a}$ on the threshold value $\hat{\mathbf{C}}$:**

$$\frac{\partial \sqrt{X}}{\partial c_\mathrm{a}} = \frac{\lambda_\mathrm{a}\,(1+M_\mathrm{a}+\lambda_\mathrm{a}\,c_\mathrm{a}-4\,\delta\,c_\mathrm{a})}{\sqrt{X}} \tag{B32}$$

$$\frac{\partial n^*_\mathrm{a}}{\partial c_\mathrm{a}} = \frac{(1+M_\mathrm{a})(1+\lambda_\mathrm{a}c_\mathrm{a}+M_\mathrm{a}-\sqrt{X})}{2\lambda_\mathrm{a}c_\mathrm{a}^2} > 0 \tag{B33}$$

$$\frac{\partial \hat{C}}{\partial c_\mathrm{a}} = \frac{2\,(1+M_\mathrm{d})}{\left(1+\lambda_\mathrm{a}\,c_\mathrm{a}+M_\mathrm{a}+\sqrt{X}\right)^2\sqrt{X}}\left[(1+M_\mathrm{d})\,\sqrt{X}+(1+M_\mathrm{a})(1+M_\mathrm{a}+\lambda_\mathrm{a}\,c_\mathrm{a})\right] > 0 \tag{B34}$$

Therefore, increased $c_\mathrm{a}$ increases the invasibility area.

***Calculation:***

$$\hat{C} = \frac{2\,(1+M_\mathrm{d})\,c_\mathrm{a}}{1+\lambda_\mathrm{a}\,c_\mathrm{a}+M_\mathrm{a}+\sqrt{X}} \tag{B35}$$

With:

$$X = (1+M_\mathrm{a}-\lambda_\mathrm{a}\,c_\mathrm{a})^2+4\,\lambda_\mathrm{a}\,c_\mathrm{a}\,\left(1+M_\mathrm{a}-\delta\,c_\mathrm{a}\right) \tag{B36}$$

$$\frac{\partial \hat{C}}{\partial c_\mathrm{a}} = \frac{1}{\left(1+\lambda_\mathrm{a}\,c_\mathrm{a}+M_\mathrm{a}+\sqrt{X}\right)^2}\left[2\,(1+M_\mathrm{d})\,\left(1+\lambda_\mathrm{a}\,c_\mathrm{a}+M_\mathrm{a}+\sqrt{X}\right)-2\,(1+M_\mathrm{d})\,c_\mathrm{a}\,\left(\lambda_\mathrm{a}+\frac{\partial \sqrt{X}}{\partial c_\mathrm{a}}\right)\right] \tag{B37}$$

$$\frac{\partial \hat{C}}{\partial c_\mathrm{a}} = \frac{1}{\left(1+\lambda_\mathrm{a}\,c_\mathrm{a}+M_\mathrm{a}+\sqrt{X}\right)^2}\left[2\,(1+M_\mathrm{d})\,\left(1+\lambda_\mathrm{a}\,c_\mathrm{a}+M_\mathrm{a}+\sqrt{X}\right)-2\,(1+M_\mathrm{d})\,c_\mathrm{a}\,\left(\lambda_\mathrm{a}+\frac{\lambda_\mathrm{a}\,(1+M_\mathrm{a}+\lambda_\mathrm{a}\,c_\mathrm{a}-4\,\delta\,c_\mathrm{a})}{\sqrt{X}}\right)\right] \tag{B38}$$

$$\frac{\partial \hat{C}}{\partial c_\mathrm{a}} = \frac{2\,(1+M_\mathrm{d})}{\left(1+\lambda_\mathrm{a}\,c_\mathrm{a}+M_\mathrm{a}+\sqrt{X}\right)^2}\left[1+\lambda_\mathrm{a}\,c_\mathrm{a}+M_\mathrm{a}+\sqrt{X}-c_\mathrm{a}\,\left(\lambda_\mathrm{a}+\frac{\lambda_\mathrm{a}\,(1+M_\mathrm{a}+\lambda_\mathrm{a}\,c_\mathrm{a}-4\,\delta\,c_\mathrm{a})}{\sqrt{X}}\right)\right] \tag{B39}$$

$$\frac{\partial \hat{C}}{\partial c_\mathrm{a}} = \frac{2\,(1+M_\mathrm{d})}{\left(1+\lambda_\mathrm{a}\,c_\mathrm{a}+M_\mathrm{a}+\sqrt{X}\right)^2}\left[1+M_\mathrm{a}+\sqrt{X}-\frac{\lambda_\mathrm{a}\,c_\mathrm{a}\,(1+M_\mathrm{a}+\lambda_\mathrm{a}\,c_\mathrm{a}-4\,\delta\,c_\mathrm{a})}{\sqrt{X}}\right] \tag{B40}$$

$$\frac{\partial \hat{C}}{\partial c_{\mathrm{a}}} = \frac{2\,(1+M_{\mathrm{d}})}{\left(1+\lambda_{\mathrm{a}}\,c_{\mathrm{a}}+M_{\mathrm{a}}+\sqrt{X}\right)^2\sqrt{X}}\left[(1+M_{\mathrm{a}})\sqrt{X}+X-\lambda_{\mathrm{a}}\,c_{\mathrm{a}}\,(1+M_{\mathrm{a}}+\lambda_{\mathrm{a}}\,c_{\mathrm{a}}-4\,\delta\,c_{\mathrm{a}})\right] \tag{B41}$$

$$\frac{\partial \hat{C}}{\partial c_{\mathrm{a}}} = \frac{2\,(1+M_{\mathrm{d}})}{\left(1+\lambda_{\mathrm{a}}\,c_{\mathrm{a}}+M_{\mathrm{a}}+\sqrt{X}\right)^2\sqrt{X}}\left[(1+M_{\mathrm{a}})\sqrt{X}+(1+M_{\mathrm{a}})^2+\lambda_{\mathrm{a}}\,c_{\mathrm{a}}\,(1+M_{\mathrm{a}})\right] \tag{B42}$$

$$\frac{\partial \hat{C}}{\partial c_{\mathrm{a}}} = \frac{2\,(1+M_{\mathrm{d}})}{\left(1+\lambda_{\mathrm{a}}\,c_{\mathrm{a}}+M_{\mathrm{a}}+\sqrt{X}\right)^2\sqrt{X}}\left[(1+M_{\mathrm{a}})\sqrt{X}+(1+M_{\mathrm{a}})(1+M_{\mathrm{a}}+\lambda_{\mathrm{a}}\,c_{\mathrm{a}})\right] \tag{B43}$$
