## Supplementary figures for "Uncovering the effects of Müllerian mimicry on the evolution of conspicuousness in colour patterns"

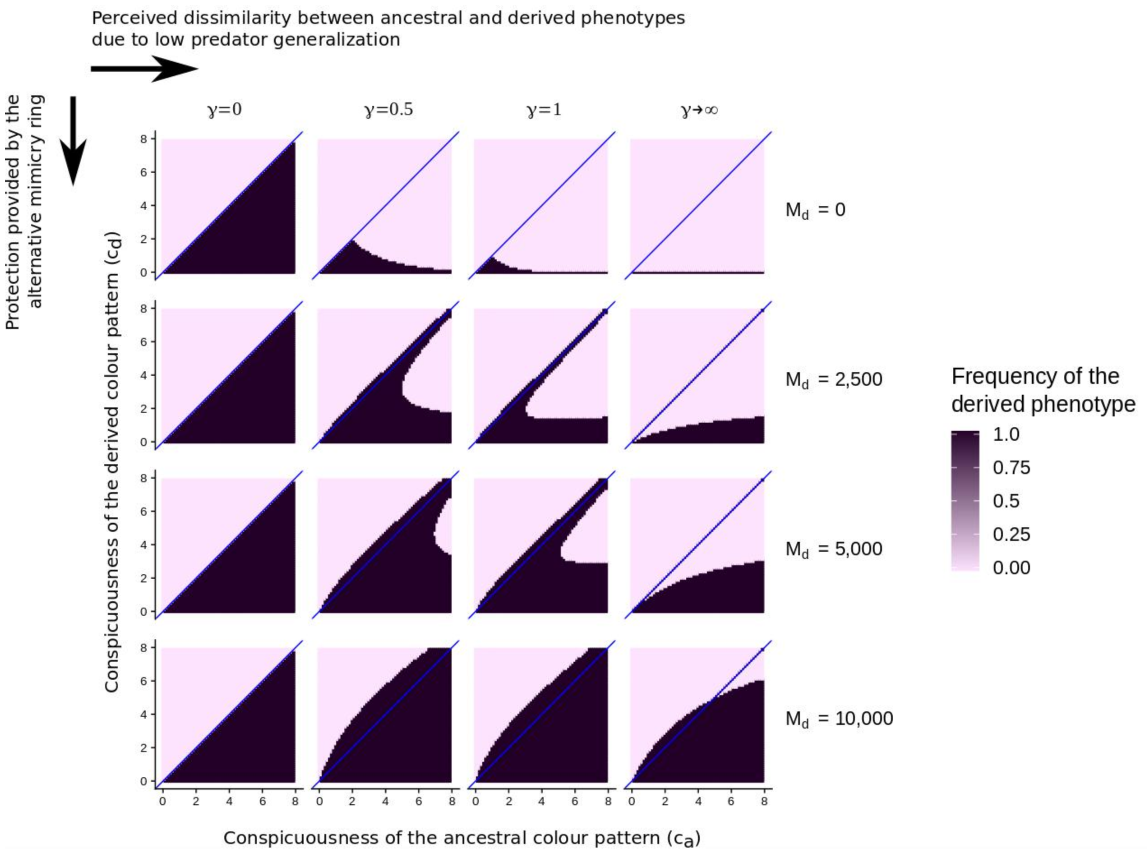


**Supplementary Figure 1. Numerical simulations showing that the derived phenotype gets fixed once it has invaded the population.** We consider the same parameter values as in Figure 2. This time, we show the frequency of mutant colour patterns after a runtime equal to $10^{20}$ using a numerical resolution method (Runge Kutta 4). Note the absence of intermediate shades of grey in all panels. When the derived phenotype invades (as shown in Figure 2 in black and red), the derived phenotype ultimately replaces the ancestral phenotype, and reaches a frequency equal to 1 (dark purple here). By contrast, when the derived phenotype does not invade (as shown in Figure 2 in grey), the derived phenotype ultimately reaches a frequency equal to 0 (light purple here).

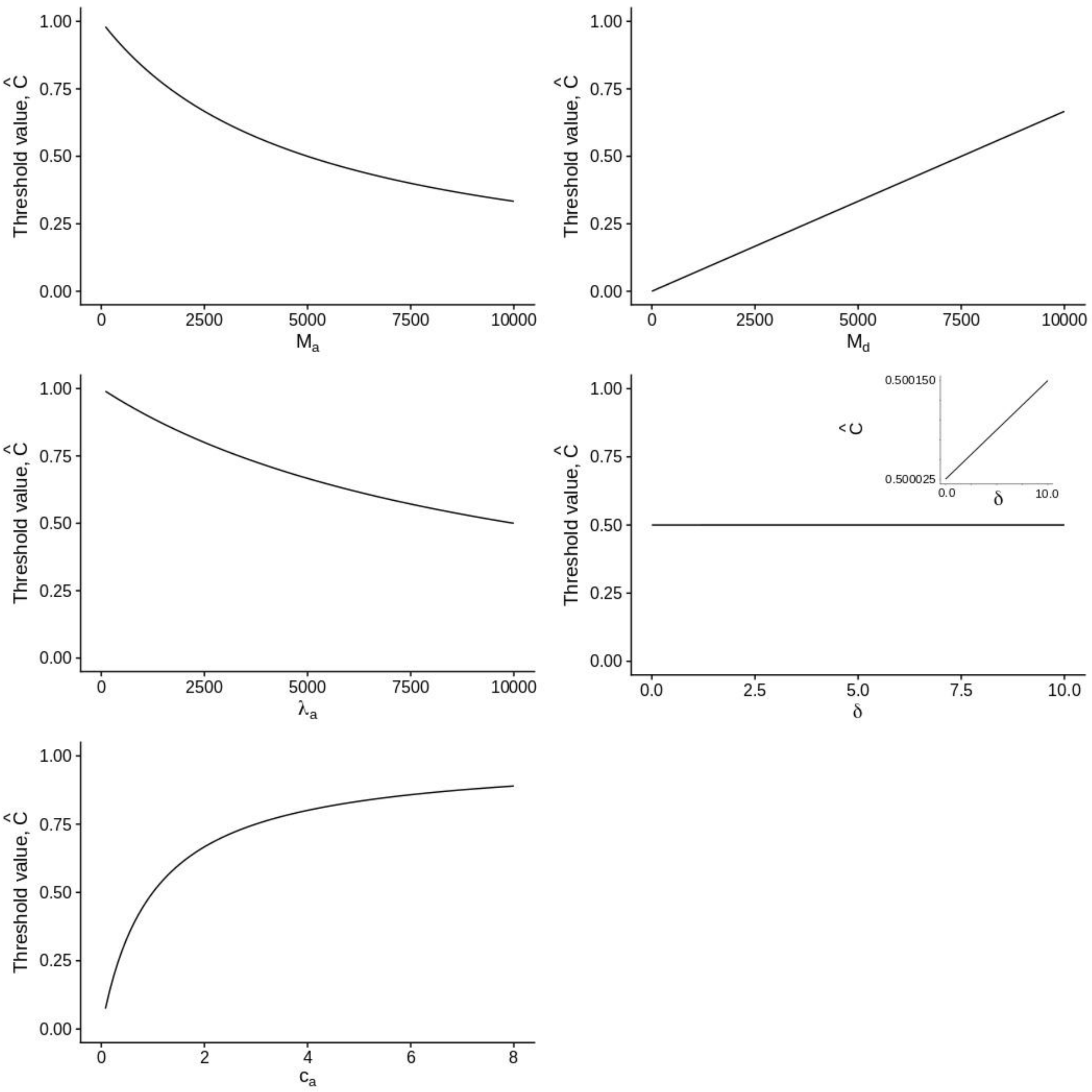


**Supplementary Figure 2. Effect of the model parameters on the threshold value below which conspicuousness is favoured (illustrating the results shown in Tab. 2).** The derived phenotype can invade when its conspicuousness $c_d$ is lower than the conspicuousness threshold $\widehat{C} = c_a \frac{1+M_d}{1+\lambda_a c_a n_a^* + M_a}$ (see Eq. [10]). Here, we represent in each subfigure how variations in a given parameter affect this threshold value. If the increase of a parameter value (e.g., increased $c_a$) increases $\widehat{C}$, this means that the range of conspicuousness values enabling the invasion of the derived phenotype increases. In the case of variations in δ, the inset shows the change in $\widehat{C}$ which is not visible by using the common y-scale ranging from 0 to 1. See default values in Table 1 (remembler that $\delta = \frac{p}{r}$ is the rescaled baseline mortality rate; and $\lambda_a = uK\beta_a$ is the rescaled deterrence factor). Note that $\lambda_\mathrm{d} = uK\beta_\mathrm{d}$ has no effect on $\widehat{C}$ (not shown here).

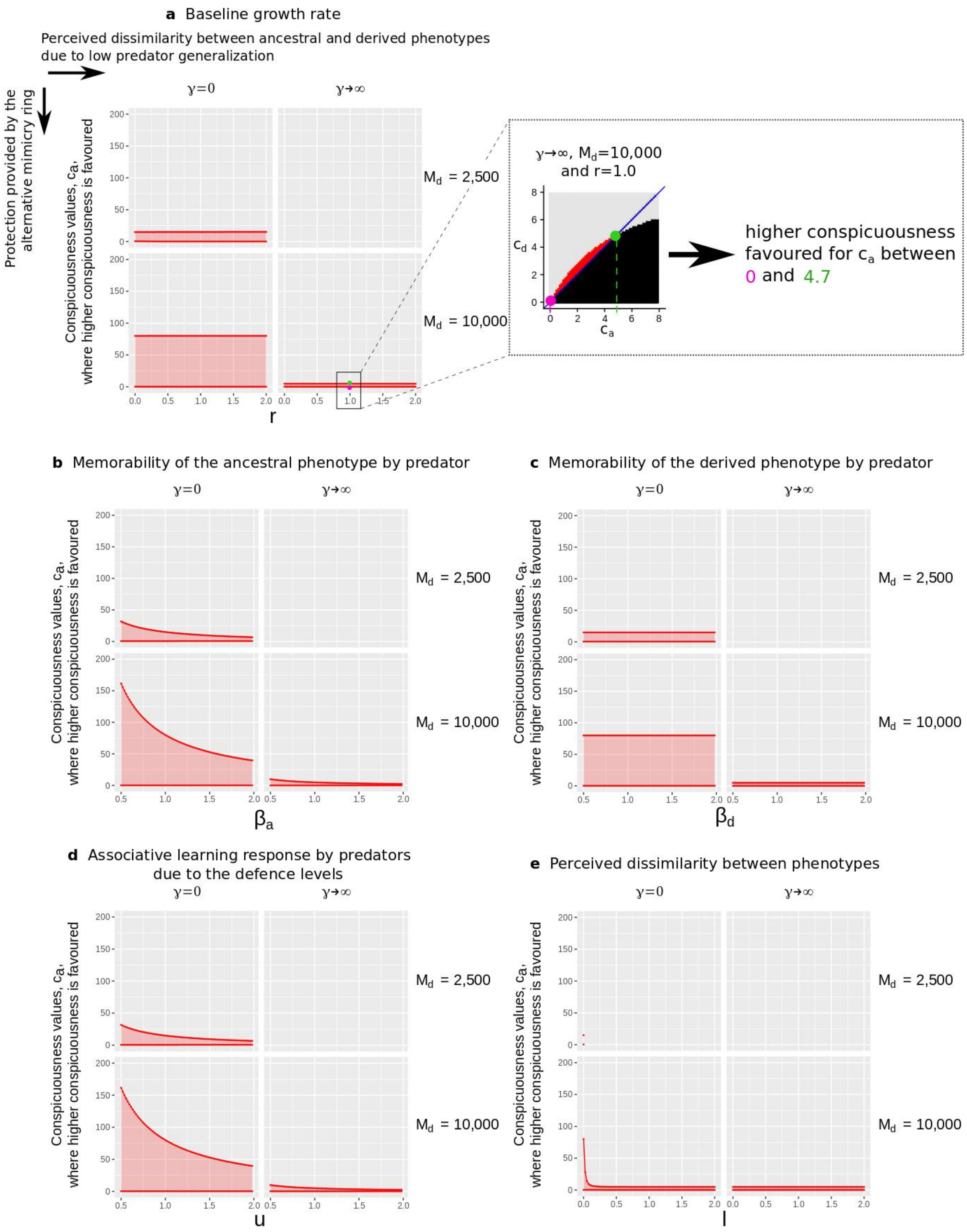


**Supplementary Figure 3. Conditions under which higher conspicuousness is favoured.** In each of the five subfigures, we vary the values of parameters $\gamma$ and $M_d$ in the four panels. Additionally, the x-axis represents the variation of different parameters in each subfigure: $r$ (a), $\beta_a$ (b), $\beta_d$ (c), $u$ (d) and $l$ (e). On the y-axis, we show the conspicuousness values, $c_a$, where higher conspicuousness is favoured. In subfigure a, we represent for a combination of parameters what these minimum and maximum values of $c_a$ correspond to in the graphs presented in the main manuscript. See default values in Table 1.

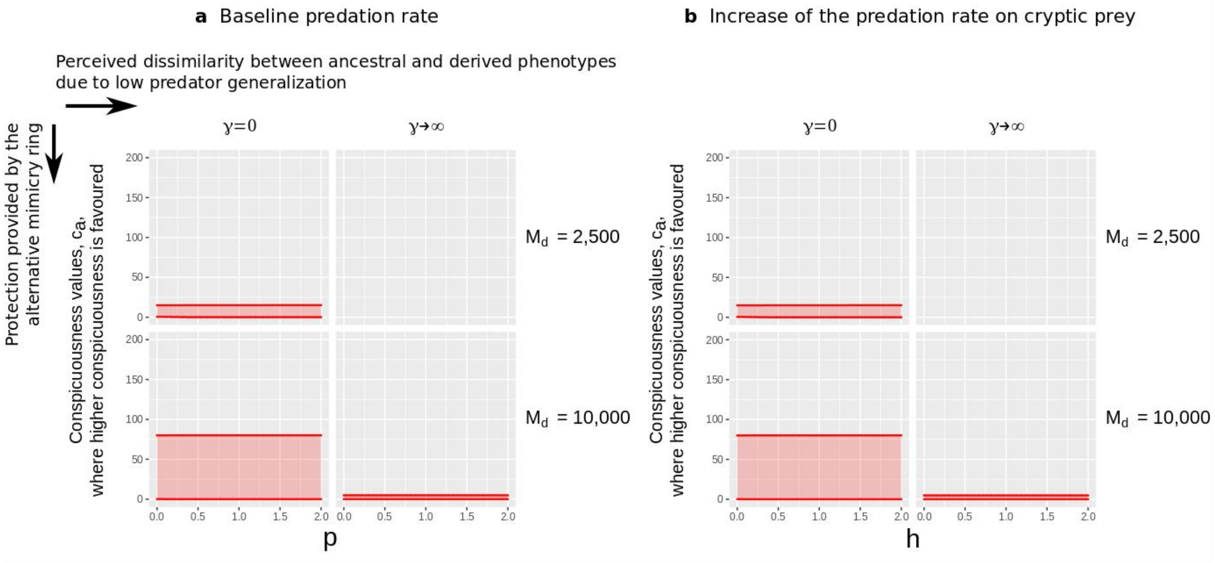


**Supplementary Figure 4. Conditions under which higher conspicuousness is favoured for various predation rates.** Same as Supp. Fig. 3 but here in each subfigure, the x-axis represents variations of $p$ (a) and $h$ (b). See Supp. Fig. 3 for more details. See default values in Table 1.

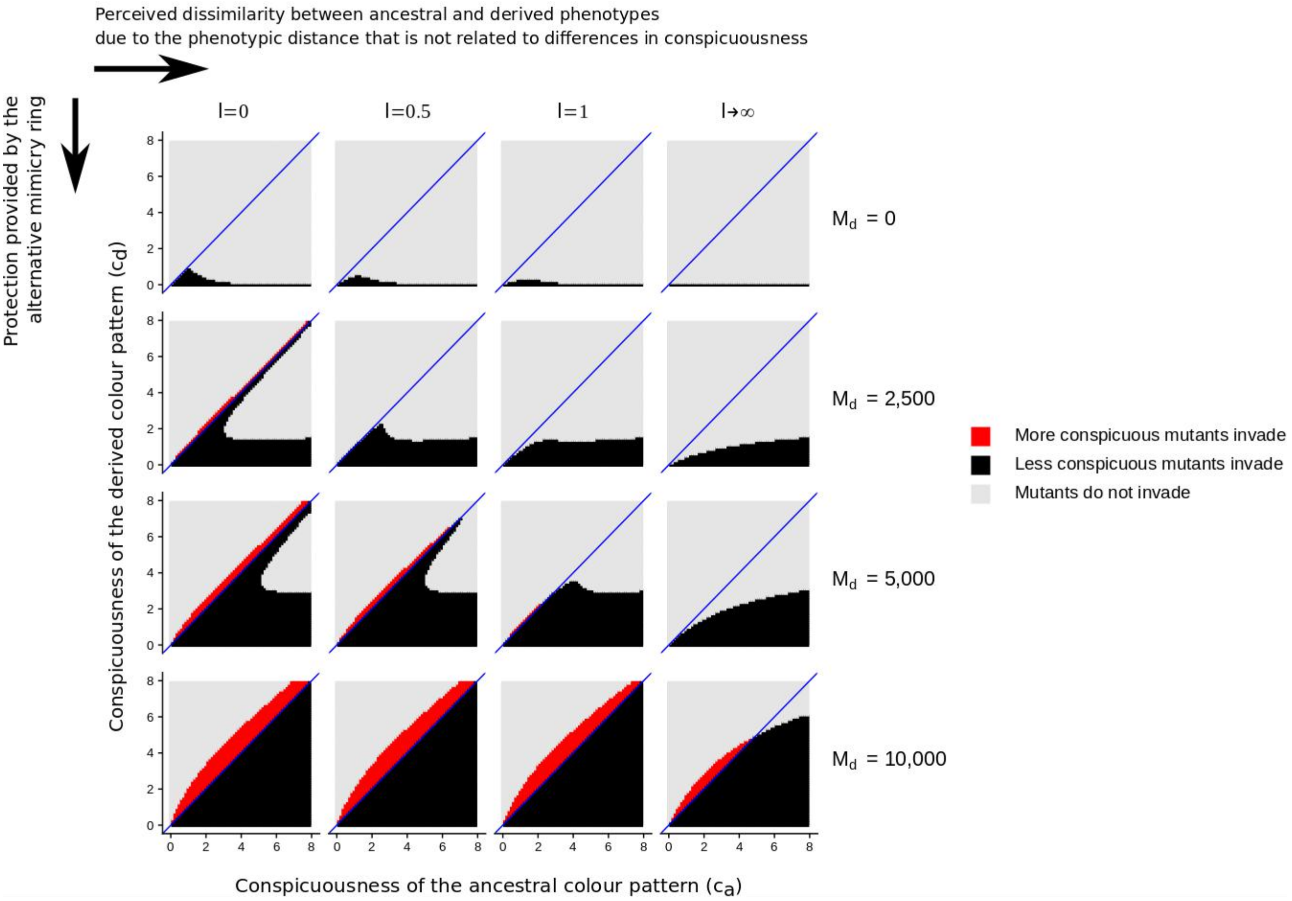


**Supplementary Figure 5. Evolution of conspicuousness depending on differences other than in conspicuousness.** We consider different values of parameter l, which controls the phenotypic distance unrelated to conspicuousness. The condition $l \rightarrow \infty$ is obtained by setting $S = 0$, just like for $\gamma \rightarrow \infty$ in Figure 2. When ancestral and derived phenotypes are different ($l \rightarrow \infty$), it is difficult for more conspicuous mutants to invade the population because the ancestral phenotype benefits from a greater number-dependent protection. By contrast, if the ancestral and derived phenotypes are totally similar ($l = 0$) or imperfect mimics (intermediate l), more conspicuous mutants are favoured because they benefit from an increased number-dependent protection by resembling both the wild-type and mutant mimetic community. See Figure 2 for more details. Here, $\gamma = 1$, $\beta_a = 1$, and $h = 0$. See other default values in Table 1.

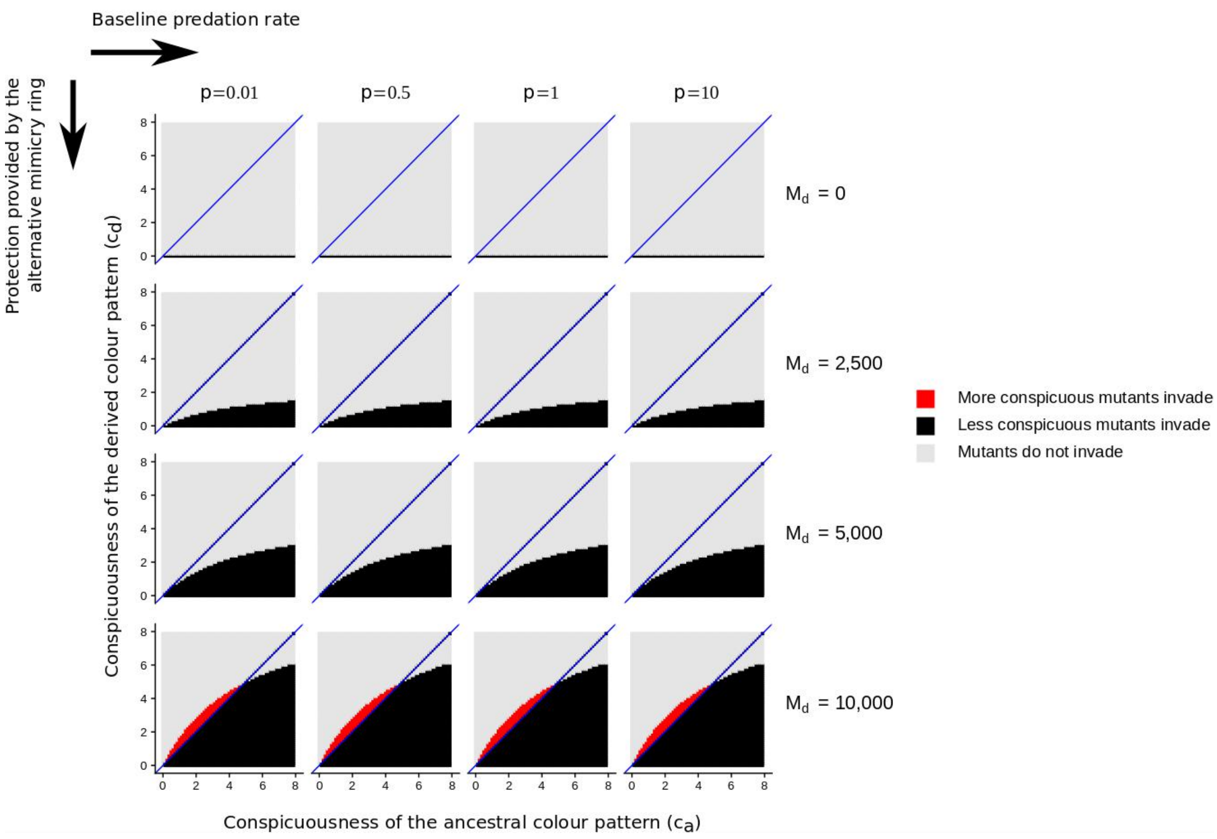


**Supplementary Figure 6. Evolution of conspicuousness depending on the baseline predation rate.** We consider different values of parameter $p$, which controls the baseline predation rate. The baseline predation rate has very little effect on the conditions of invasion of the derived phenotype. See Figure 2 for more details. Here, $\gamma \rightarrow \infty$, $\beta_a = 1$, and $h = 0$. See other default values in Table 1.
