## Supplementary tables for "Uncovering the effects of Müllerian mimicry on the evolution of conspicuousness in colour patterns"

### SUPPLEMENTARY TABLE

**Supplementary Table 1. Sensitivity of the invasion of the derived phenotypes to the different parameters, assuming a ‘complete mimicry shift’.** The derived phenotype can invade when its conspicuousness $c_d$ is lower than the conspicuousness threshold $\widehat{C} = c_a \frac{1+M_d}{1+\lambda_a c_a n_a^* + M_a}$ (see Eq. [10] and Appendix B). See also Supp. Fig. 2.

| Parameter | Sensitivity of the invasion condition | Meaning |
|---|---|---|
| $M_a$, the protection brought by the *ancestral* mimicry ring | $\frac{\partial \widehat{C}}{\partial M_a} < 0$ | Higher protection provided by the mimicry ring matching the ancestral phenotype ($M_a$) **decreases** the range of conspicuousness values enabling the invasion of the derived phenotype. |
| $M_d$, the protection brought by the *derived* mimicry ring | $\frac{\partial \widehat{C}}{\partial M_d} > 0$ | Higher protection provided by the mimicry ring matching the derived phenotype ($M_d$) **increases** the range of conspicuousness values enabling the invasion of the derived phenotype. |
| $\lambda_a$, the rescaled deterrence factor associated with the ancestral phenotype ($\lambda_a = uK\beta_a$) | $\frac{\partial \widehat{C}}{\partial \lambda_a} < 0$ | Higher deterrence factor $\lambda$ **decreases** the range of conspicuousness values enabling the invasion of the derived phenotype. |

| | | |
|---|---|---|
| $\delta$, rescaled baseline mortality rate ($\delta = p/r$) | $\frac{\partial \hat{C}}{\partial \delta} > 0$ | Increased baseline predation pressure ($\delta$) **increases** the range of conspicuousness values enabling the invasion of the derived phenotype. |
| $c_a$, conspicuousness of the ancestral phenotype | $\frac{\partial \hat{C}}{\partial c_a} > 0$ | Higher conspicuousness of the ancestral phenotype ($c_a$) **increases** the range of conspicuousness values enabling the invasion of the derived phenotype. |
